# AAV NRF2 Gene Therapy Preserves Retinal Structure and Function in Rodent Models of Oxidative Damage

**DOI:** 10.1101/2025.07.09.663952

**Authors:** Apolonia Gardner, Shuai Wang, Adam Daniels, Dan Li, Christine Wu, Lucas Lin, Christin Hong, Sophia R. Zhao, Kamil Kruczek, Genevieve Weist, Laura Barrio Real, Virginia Haurigot, Richard T. Born, Constance L. Cepko

**Author notes:** Correspondence should be addressed to C.L.C., Department of Genetics, Harvard Medical School, 77 Avenue Louis Pasteur, Boston MA 02115, USA, Ph. Denotes co-first authors (Apolonia Gardner, Shuai Wang).

## Abstract

Dry age-related macular degeneration is the most frequent cause of visual impairment in individuals over age 50 in developed countries. It is characterized by subretinal deposits of oxidized proteins and lipids and results in progressive loss of high acuity vision. One major risk factor is smoking, which causes oxidative stress in many tissues, including the eye. We previously showed that an adeno-associated viral vector expressing human NRF2 (AAV8/Best1-Nrf2), a transcription factor that regulates responses to oxidative damage, slowed degeneration in mouse models of another blinding disorder, retinitis pigmentosa, which also includes oxidative stress. Here, our AAV8/Best1-Nrf2 vector was tested in a model of oxidative stress wherein sodium iodate was injected systemically, as this is often used to model dry age-related macular degeneration. Sodium iodate causes acute oxidative damage to supporting cells of the retina, the retinal pigment epithelial cells, and ultimately leads to photoreceptor death. Subretinal injection of AAV8/Best1-Nrf2 led to protection of the retinal pigment epithelium and photoreceptors, as well as preservation of visual function, in rat and mouse sodium iodate models. AAV8/Best1-Nrf2 may serve as an effective gene-agnostic therapy for diseases with oxidative stress, including dry age-related macular degeneration.

**ETOC:** Oxidative stress occurs in diseases that lead to vision loss, including age-related macular degeneration (AMD). AMD is commonly modeled by injection of sodium iodate (S.I.), an oxidizing agent. AAV-mediated delivery of Nrf2, a transcription factor that regulates oxidative stress, protects the eye from S.I. damage in rat and mouse models.

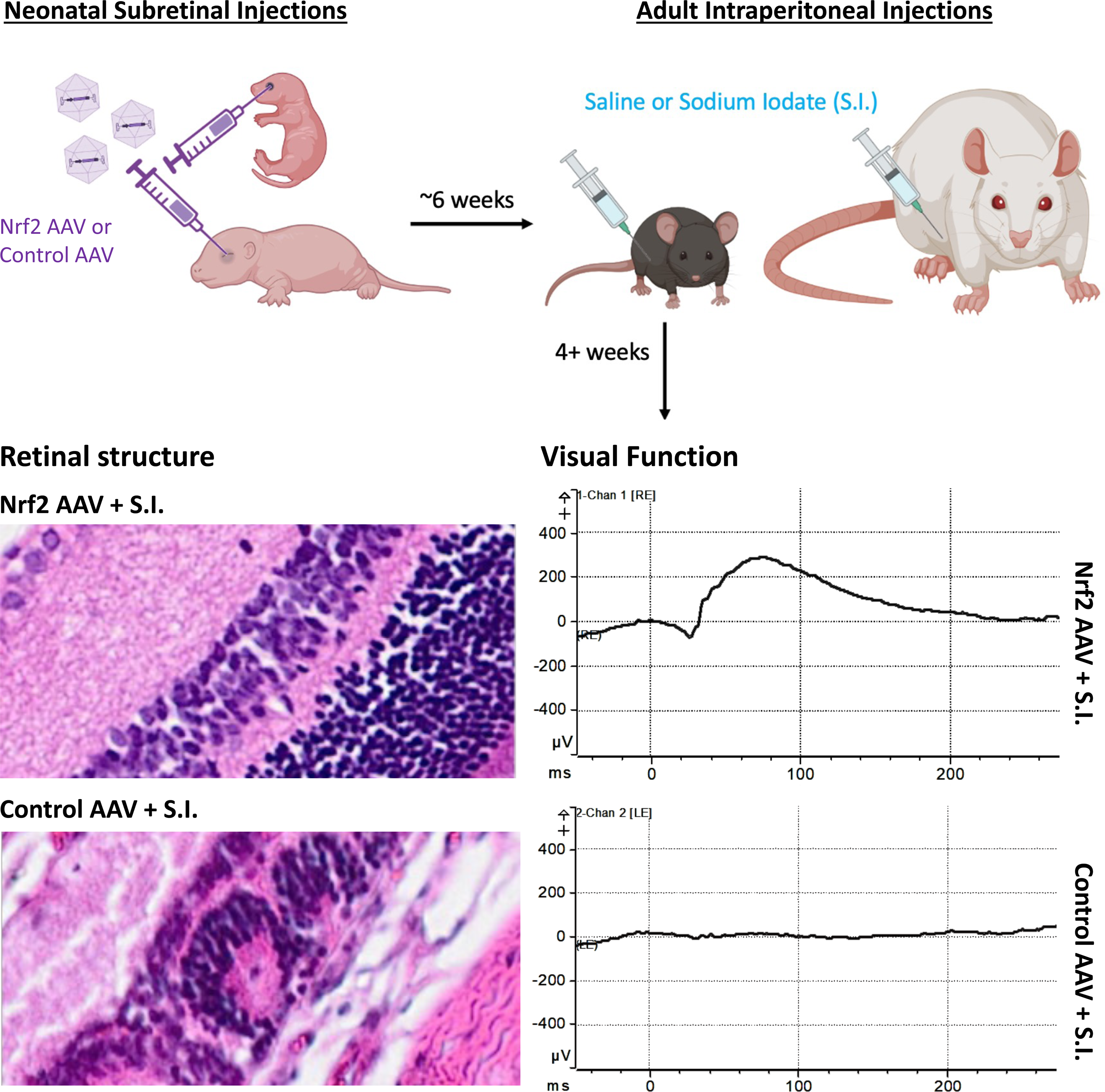

## Introduction

Age-related macular degeneration (AMD) is a disease leading to loss of high acuity vision, primarily in individuals over 50 years of age.^1^ There are two forms of AMD, dry and wet, where dry AMD comprises ∼90% of all AMD cases.^2–5^ A hallmark symptom of AMD is the appearance of drusen, subretinal deposits containing oxidized proteins and lipids.^6^ These form preferentially in the macula, an area of the retina surrounding the fovea, which is responsible for high acuity vision.^7,8^ The disease may originate, however, from systemic problems that initially affect a retina-adjacent supporting cell monolayer, the retinal pigment epithelium (RPE).^9^ RPE cells have several roles, such as providing nutrients to photoreceptors and removing their waste, e.g., by phagocytosing photoreceptor outer segments which are routinely shed.^10^ RPE loss in patients with dry AMD generally transitions from small to progressively larger patches over many years. Geographic atrophy (GA) occurs when substantial areas of the RPE are lost in advanced stages of the disease.^11,12^ Wet AMD, a less common and more severe form of AMD, can occur in advanced stages of dry AMD or arise independently.^8^ Unlike dry AMD, “wet” AMD is so named because vasculature in the eye typically enlarges and hemorrhages, damaging the retina and obstructing vision.^3^ FDA-approved angiogenesis inhibitors (e.g., VEGF inhibitors^2^) are available and effective for wet AMD, and there are two FDA-approved complement inhibitors (pegcetacoplan and avacincaptad) to treat GA associated with dry AMD.^13^ Despite reducing the rate of GA expansion over time, a lack of improvement in vision was observed in clinical trials 1 or 2 years after treatment with complement inhibitors.^14^ Nutritional supplements that contain antioxidants, as in the AREDS2 formulation, provide a different approach for dry AMD.^15^ In the AREDS2 clinical trial, patients showed a ∼25% reduction in the risk of progression from intermediate to advanced dry AMD after taking this supplement.^16^ Albeit promising, the ocular potency of nutritional supplementation is low, leaving room for additional treatments to improve patient outcomes.

Many age-related diseases, including AMD, are thought to be caused, at least in part, by oxidative stress-induced damage.^17,18^ The eye is thought to be particularly susceptible to oxidative damage due to the high oxygen demands of photoreceptors and the high light environment.^19^ We previously showed that delivery of a transcription factor, NRF2, reduced the loss of cone photoreceptors, protected the RPE, and prolonged vision in mouse models of another retinal degenerative disease, retinitis pigmentosa.^20,21^ This therapy was effective in 3 mouse models of retinitis pigmentosa (3 different genotypes), potentially making it a disease gene-agnostic therapy. NRF2 delivery was achieved using adeno-associated virus (AAV), a gene therapy vector that is the vector of choice for ocular diseases.^22^ AAV vectors are favorable for in vivo applications due to their nonpathogenicity, broad tissue tropism, high titer, and stable episomal genome in non-dividing cells.^23^ In addition, AAV vectors can be engineered to express a gene specifically in the RPE or photoreceptors.

Nrf2 is a transcription factor that undergoes proteasomal degradation in the cytoplasm under homeostatic conditions, but upon oxidative or xenobiotic stress, translocates to the nucleus and activates target genes to combat the effects of oxidative stress.^20,21,24,25^ A number of studies have tested the ability of Nrf2 to reduce stress-related phenotypes or pathological symptoms in genetic diseases.^26^ Nrf2 levels can be raised by targeting its interaction with Keap1,^27^ the protein that directs its ubiquitination and proteasomal degradation, e.g. by mutating amino acids on Nrf2 that mediate the Keap1 interaction,^28^ or by genetic deletion of Keap1.^29^ There are also dietary compounds touted to be beneficial in aging and a variety of diseases, with effects thought to be mediated by Nrf2.^30,31^ Treatment with a small molecule, such as dimethyl fumarate, as is done for multiple sclerosis, is also thought to increase NRF2 activity.^32^ In addition to fighting oxidative damage, Nrf2 might help dampen inflammation. Interactions with other transcription factors were shown to repress expression of Il6 and Il1b,^33^ two cytokines often upregulated when innate immunity is activated, or in early stages of adaptive immunity. NRF2 was thus chosen as a potential agent to reduce oxidative damage and inflammation in mouse models of retinitis pigmentosa.^20,21^ The success of this treatment in retinitis pigmentosa suggested it might likewise relieve oxidative stress and inflammation in other diseases, such as dry AMD.

Several models of dry AMD exist, each with their own limitations,^34^ but one of the most commonly used models of dry AMD is rooted in oxidative stress caused by sodium iodate (S.I.).^35,36^ S.I. is an oxidizing chemical that, when administered through various routes, rapidly causes RPE cell death. RPE loss occurs in patches, with loss occurring preferentially in the central area,^37^ much as AMD occurs in the centrally located macula in humans. Injection of S.I. is simple, reproducible, and is often used in tests of dry AMD treatments. Here, this model was used to test whether AAV8/Best1-Nrf2 could protect against S.I.-induced damage. The expression of NRF2 was limited to the RPE using the BEST1 promoter,^21,38^ as this tissue is affected in AMD and might also be the origin of damage in AMD.^1^ A combination of visual function tests (electroretinography, or ERG), imaging (optical coherence tomography, or OCT), and histological measurements showed that AAV8/Best1-Nrf2 effectively counteracted S.I.-induced RPE and photoreceptor cell death. As well, it preserved visual function in both mouse and rat models.

## Results

### Characterization of S.I. Induced Effects in Rodent Models

An S.I. IP dose of 75 mg/kg was selected for both rats and mice guided by prior literature and preliminary experiments.^39^ Ocular tissues were assayed for RPE loss, photoreceptor cell loss, retinal thinning, and visual function.

To determine RPE cell phenotypes following S.I. or saline IP injections, sclera/choroid/RPE flatmounts were stained with phalloidin.^40^ Phalloidin binds to F-actin, effectively outlining RPE cell morphologies in the RPE monolayer.^40^ RPE flatmounts from saline injected animals showed the characteristic hexagonal, honeycomb-like morphology of healthy RPE cells, whereas S.I. administration reproducibly resulted in loss of phalloidin signal in the central ∼75% of each flatmount in rats and mice (Figure 1A and Figure S1A). Peripheral RPE cells were consistently observed to survive in all flatmounts. Quantification of RPE cells remaining in mid-peripheral areas of each flatmount (calculated as 50% of the distance between the optic nerve and outer edges of each flatmount) is shown in Figure 1D and Figure S1D. Loss of phalloidin staining in the central areas of RPE flatmounts, as well as loss of RPE65 staining in central areas of cross sections of eyecups prepared from rats administered saline or S.I. (see Methods, Figure S2A-B), suggests that RPE cells in the central regions died upon S.I. exposure.

**Figure 1:**
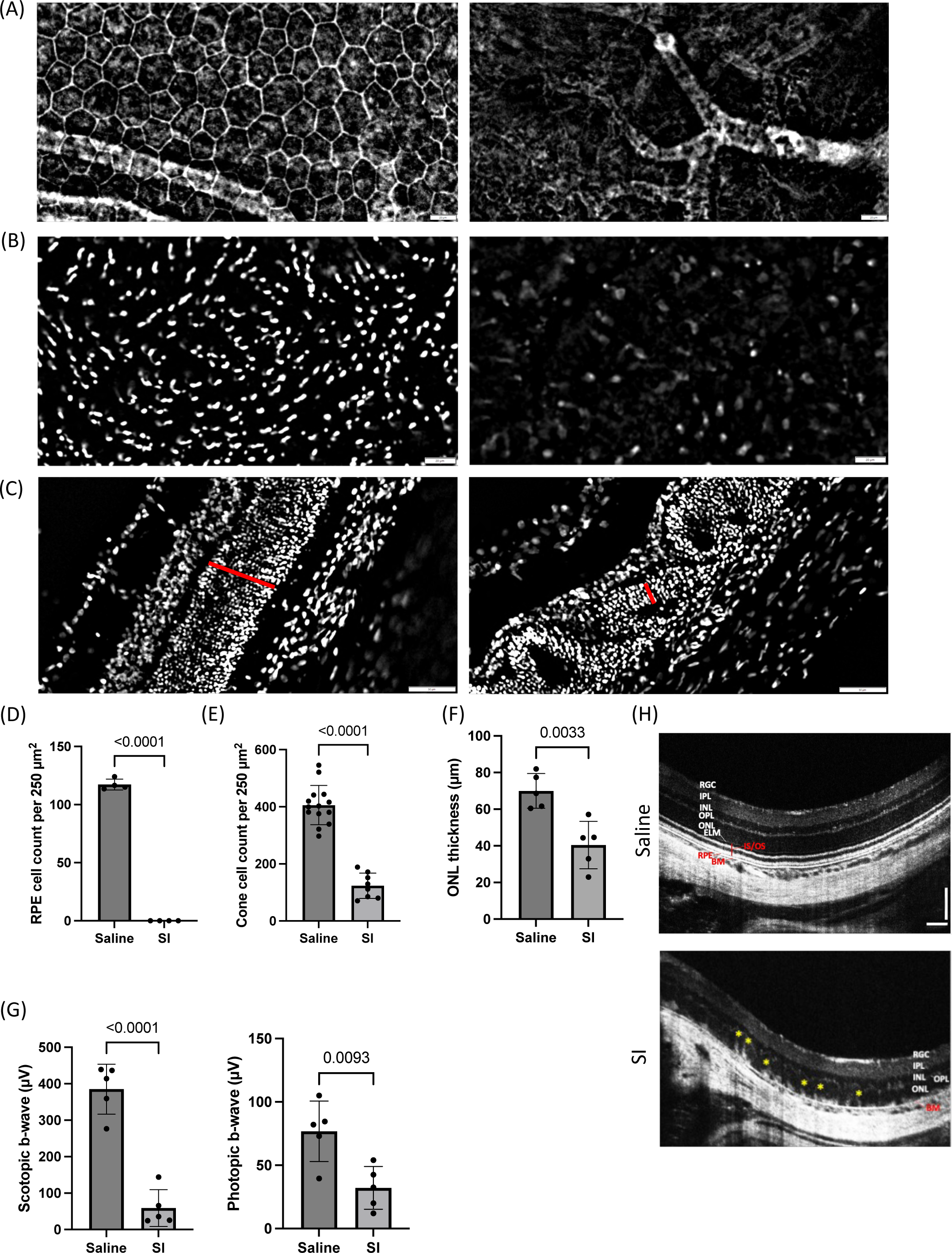
Assessment of RPE and retinal histology after IP injection of S.I. in rats. Sprague-Dawley rats were IP injected with saline or S.I. at 6-8 weeks of age. Tissue was harvested 4-5 weeks post IP injection. (A) Representative RPE flatmounts (central region) from a saline (left) or S.I. (right) IP-injected rat. Flatmounts were stained with phalloidin (white). Scale bar is 20 microns. (B) Representative retinal flatmounts (central region) from a saline (left) or S.I. (right) IP-injected rat. Flatmounts were stained with an antibody to the cone marker, CAR (white). Scale bar is 20 microns. (C) Representative eyecup sections (mid-peripheral region) from a saline (left) or S.I. (right) IP-injected rat. Sections were stained with DAPI (white). Red bar indicates the ONL. Scale bar is 50 microns. (D) Quantification of the number of RPE cells remaining (n=4 eyes per group, mean ± SD, p<0.0001; unpaired t-test). (E) Quantification of the number of cones remaining (n=14 saline and n=8 S.I. eyes, mean ± SD, p<0.0001; unpaired t-test). (F) Quantification of the average ONL thickness from eyecup sections (n=5 per group, mean ± SD, p=0.0033; unpaired t-test). (G) Scotopic ERG b-wave amplitudes (at a 0.1 cd.s/m^2^ flash stimulus, left) and photopic ERG b-wave amplitudes (at a 10 cd.s/m^2^ flash stimulus, right) for rats injected with saline or S.I. were collected 3-5 weeks post IP injection (n=5 per group, mean ± SD, p<0.0001 for scotopic and p=0.0093 for photopic; unpaired t-test for each plot). (H) Representative OCT images for a saline IP-injected rat (top panel) or an S.I. IP-injected rat (bottom panel) at 4 weeks post IP injection. Major retinal layers are annotated as follows: GCL – ganglion cell layer; IPL – inner plexiform layer; INL – inner nuclear layer; OPL – outer plexiform layer; ONL – outer nuclear layer; ECM – external limiting membrane; IS/OS – inner and outer segments of photoreceptors; RPE – retinal pigment epithelium; BM – Bruch’s membrane. Rosettes are highlighted with yellow asterisks. Scale bar is 100 microns.

To determine the effects of S.I. IP injections on cones, retinal flatmounts were isolated from the same eyes used for RPE flatmounts and were subjected to immunohistochemistry (IHC) for the cone-specific protein, cone arrestin (CAR). CAR is localized throughout the cones, including within cone outer segments,^41^ enabling surviving cones to be recognized as dots on retinal flatmounts. After imaging entire retinal flatmounts, automated image processing algorithms were applied (see Methods) to count the number of dots in mid-peripheral regions of the flatmount. As with RPE, cones in the most peripheral regions of retinal flatmounts typically survived. However, cones located in the central ∼75% of each flatmount (including “mid-peripheral” cones, or cones quantified in boxes of fixed size placed halfway between the optic nerve and outer edges of the flatmount) lost CAR staining following S.I. injection. Quantification showed 4-fold fewer mid-peripheral CAR+ cones in rats and mice injected IP with S.I. compared to saline injected controls (Figure 1B/E and Figure S1B/E).

To determine photoreceptor survival post S.I. injection, the thickness of the outer nuclear layer (ONL) was measured at a fixed distance away from the optic nerve in eyecup sections (see Methods). The ONL comprises primarily rods, as they are >90% of photoreceptors in this layer.^42^ The average ONL thickness was 70 ± 9 µm for rats and 54 ± 15 µm for mice injected IP with saline, and it was reduced to 40 ± 13 µm and 26 ± 8 µm for S.I.-injected rats and mice, respectively (Figure 1C/F and Figure S1C/F).

Visual function analyses were also carried out. ERG is used to evaluate the overall physiological status of an animal’s vision^43^ and was evaluated here at baseline, just before S.I. injection at ∼6 weeks of age, as well as at later time points. ERG was used to measure low-light vision (scotopic b-wave, rod function) as well as brighter light vision (photopic b-wave, cone function). ERG revealed a significant loss of visual function in S.I.-injected rats (assayed 3-5 weeks post IP) and mice (assayed 3-4 weeks post IP) relative to saline controls (Figure 1G and Figure S1G).

Images of ocular structures were also collected using OCT.^44^ In contrast to saline-injected control animals, S.I.-injected rats and mice showed a loss of overall retinal structure, ONL thinning, and “rosette”-like toxicity phenotypes (Figure 1H and Figure S1H). OCT imaging revealed marked retinal thinning in mice compared to rats at the same dose of S.I. and at a similar timepoint post IP injection, suggesting mice may be inherently more sensitive to S.I. damage compared to rats.

### AAV8/Best1-Nrf2-Mediated Effects on Retinal Structure and Cells

An AAV vector expressing human NRF2, driven by an RPE-specific human promoter from the BEST1 gene^38^ and using capsid serotype 8 (AAV8/Best1-Nrf2), was previously shown to greatly improve the health of the RPE in a mouse model of retinitis pigmentosa.^21^ This vector was used to treat rodents injected intraperitoneally (IP) with S.I. in albino Sprague-Dawley rats and pigmented C57BL/6 mice (see Figure 2A for a schematic). As above, IP injections of saline served as a vehicle control.

**Figure 2:**
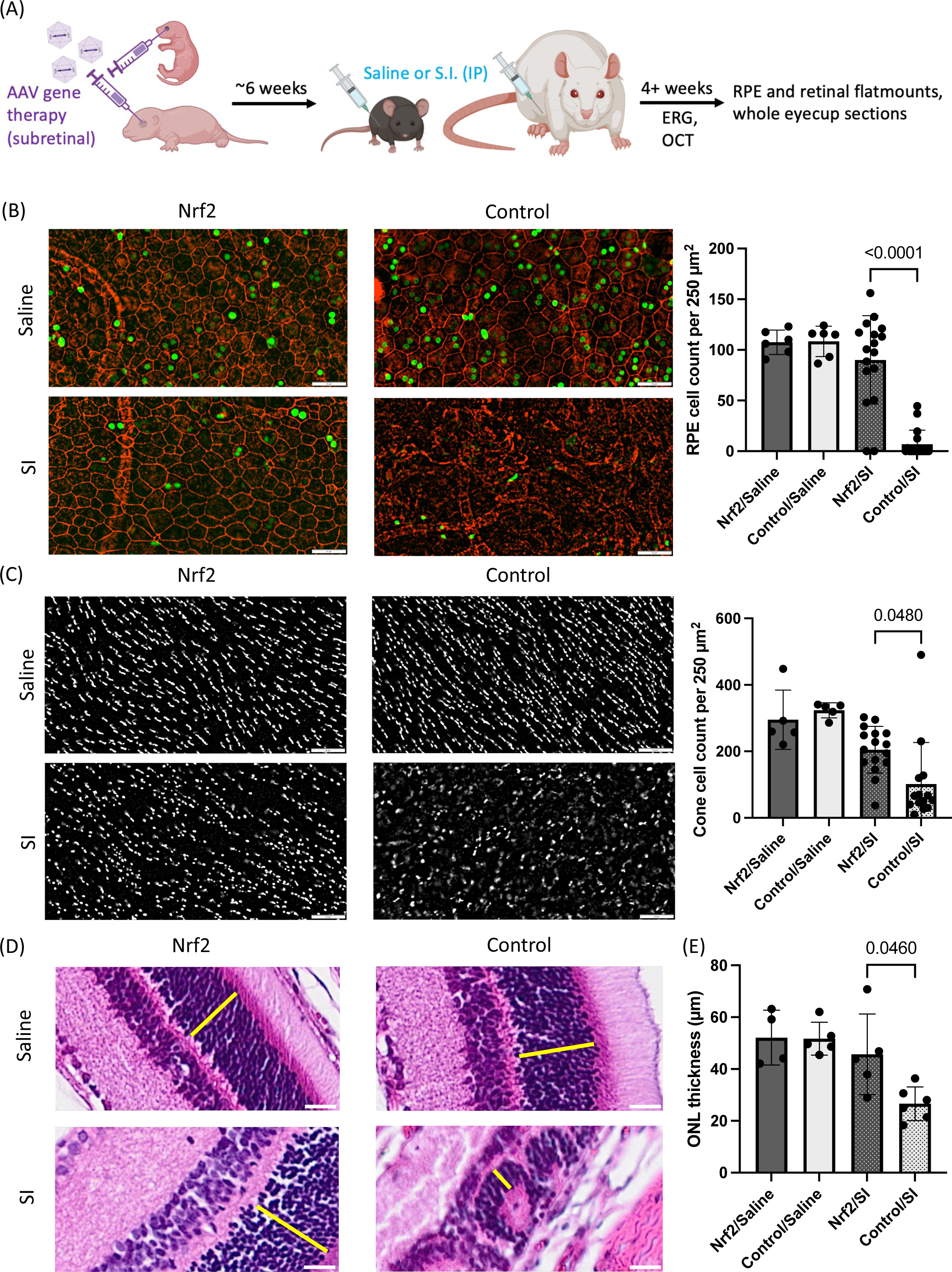
Assessment of AAV8/Best1-Nrf2 on RPE and retinal histology in rats. (A) Protocol used to assess efficacy of AAV-mediated NRF2 gene transfer in prevention of S.I induced damage. Unless otherwise noted, rats or mice were injected at birth with AAV8/Best1-Nrf2 + AAV8/RedO-H2B-GFP in one eye or AAV8/Best1-6xSTOP-mutGFP control vector + AAV8/RedO-H2B-GFP in the contralateral eye and IP injected with saline or S.I. at 6-8 weeks of age. Ocular tissues were harvested for histology at 4 or more weeks post IP injection. ERG and OCT measurements were collected at one or more timepoints just before IP injection up until just before tissue harvests. (B) Representative RPE flatmounts (central region) stained with phalloidin (red). Green dots in RPE nuclei are H2B-GFP signals, likely from AA8/RedO-H2B-GFP concatemerization with co-injected Best1 promoters to achieve RPE expression of H2B-GFP (see Results and Discussion). Scale bar is 50 microns. Quantification of the number of RPE cells is shown on the right (n=17 Nrf2/S.I. flatmounts, n=17 Control/S.I. flatmounts, n=6 Nrf2/Saline flatmounts, n=6 Control/Saline flatmounts, mean ± SD, p<0.0001; paired t-test). (C) Representative retinal flatmounts (central region) stained with an antibody to the cone marker, CAR (white). Scale bar is 50 microns. Quantification of the number of cones is shown on the right (n=16 Nrf2/S.I. flatmounts, n=14 Control/S.I. flatmounts, n=5 Nrf2/Saline flatmounts, n=5 Control/Saline eyes, mean ± SD, p=0.0480; paired t-test). (D) Rats were injected similarly to panels B-C except a 100-fold lower dose (2e7 vg) of AAV8/RedO-H2B-GFP was delivered per eye. Representative eyecup sections (mid-peripheral region) stained with hematoxylin & eosin. Yellow bar indicates the ONL. Scale bar is 50 microns. (E) Quantification of the average ONL thickness in DAPI-stained cryosections from rats injected as described in (D). (n=4-6 per injection condition, mean ± SD, p=0.0460, paired t-test).

Viral vectors were injected into the subretinal space of rat or mouse neonatal pups (see Methods for doses of vectors used in each species). As a control for the AAV injection and expression in the RPE, an equal dose of a control virus was delivered to the subretinal space of the contralateral eye. Control vectors used across all experiments were constructed so as to not have an open reading frame of >73 amino acids. The control viruses were one of the following two constructs: (1) an AAV with a BEST1 promoter, two STOP codons in tandem in each reading frame, and an EGFP sequence with 41 point mutations to eliminate any potential start codons (ATG, GTG, and TTG), called AAV8/Best1-6xSTOP-mutGFP, or (2) an AAV with a BEST1 promoter, two STOP codons in tandem in each reading frame, and no EGFP sequence, called AAV8/Best1-6xSTOP.^45^ Unless otherwise noted, another AAV, AAV8/RedO-H2B-GFP, was also co-injected into every eye to label the transduced area of each flatmount (see Methods for dosing of GFP vectors). Despite the use of the human red opsin (RedO) cone promoter for expression of H2B-GFP, the RPE was observed to express H2B-GFP. This was not due to a lack of specificity of the RedO promoter, but likely due to concatenation or recombination of the co-injected genomes (See Discussion).

Two litters of rats were injected at birth with the aforementioned vectors. To avoid bias in injection coverage in either the left or right eyes, one litter received AAV8/Best1-Nrf2 in the left eye and one litter received it in the right eye. At ∼6 weeks of age, they were injected IP with S.I. or saline. At 4-6 weeks post S.I. injection, animals were sacrificed and histology was carried out on RPE and retinal flatmounts. Rat RPE flatmounts revealed nearly complete loss of central RPE cells following injection of S.I. in animals injected with AAV8/Best1-6xSTOP-mutGFP (Figure 2B). In contrast, there was nearly complete preservation of RPE cell numbers after injection of AAV8/Best1-Nrf2, when compared with saline-injected control animals.

Cones also showed a statistically significant benefit in AAV8/Best1-Nrf2 injected eyes relative to AAV8/Best1-6xSTOP-mutGFP injected eyes, as assessed by CAR labeling of cones on flatmounts (Figure 2C). However, CAR+ cone numbers were not quite at the level of saline-injected controls. To determine if cones were indeed dying, as opposed to losing CAR as a marker, cones were counted using the RedO-H2B-GFP transduction to mark cone nuclei. Using this approach, fewer GFP+ cone nuclei were observed in animals injected with control AAV and S.I., relative to Nrf2 AAV/SI administration, but cone preservation by Nrf2 was not quite statistically significant using this metric (Figure S2C). Previous experience with this allele of GFP, which is incorporated into chromatin, has suggested that cells that have died, but have not been cleared, can retain detectable nuclear GFP (Krause and Cepko, unpublished), which might be the case here.

As a measure of rod survival, ONL thickness and retinal morphology were examined. ONL thickness was better preserved after S.I. injection in AAV8/Best1-Nrf2 injected animals compared to AAV8/Best1-6xSTOP-mutGFP injected animals. Hematoxylin & eosin (H&E) staining of sections from paraffin-embedded tissue showed rosettes and disrupted ONL in control virus-injected eyes after S.I. injection (Figure 2D). Quantification of ONL thickness on separate samples processed as cryosections stained with DAPI (i.e., not H&E, see Methods) showed a significant reduction in control virus-injected eyes relative to AAV8/Best1-Nrf2 injected eyes (Figure 2E).

Mice were subjected to a similar protocol as rats, with neonatal AAV subretinal injections and adult IP injections of S.I. Comparisons were made between eyes injected with AAV8/Best1-Nrf2 and control virus. Mice showed similar trends as rats in terms of preservation of RPE (Figure S3A/D), cones (Figure S3B/E), and ONL thickness (Figure S3C/F) by AAV8/Best1-Nrf2 after IP injection of S.I.

### *In vivo* Assessment of the Effects of AAV8/Best1-Nrf2

In addition to the *post mortem* histological assays described above, the same rats and mice were tested for retinal function and structure using ERG and OCT. Up to 3 timepoints were assayed: (1) immediately before the S.I. injection to establish baseline visual function/structure, (2) immediately before endpoint tissue harvests, and, in most cases, also (3) ∼1-2 weeks post S.I. injection, as an additional mid-experiment readout of NRF2 effects. ERGs of rats collected at baseline demonstrated healthy photopic and scotopic vision across all eyes of all rats (Figure 3A and S4). At 2 or 4 weeks post S.I. injection, linear mixed-effects models^46,47^ applied across multiple photopic ERG flash intensities showed significant preservation of photopic vision in AAV8/Best1-Nrf2-treated eyes relative to control AAV injected eyes (Figure 3B-C, Supplemental Methods). Likewise, paired t-tests at a single scotopic flash intensity demonstrated significant preservation of scotopic vision in AAV8/Best1-Nrf2-treated eyes relative to control AAV injected eyes (Figure 3B-C). Across all timepoints, saline control rats retained healthy scotopic and photopic vision regardless of whether the eye was injected with AAV8/Best1-Nrf2 or control AAV (Figure 3A-C and Figure S4). Imaging using OCT showed a retinal “rosette” toxicity phenotype in eyes of animals injected with S.I. and a control AAV, but not in eyes of animals injected with S.I. and AAV8/Best1-Nrf2. (Figure 3D).

**Figure 3:**
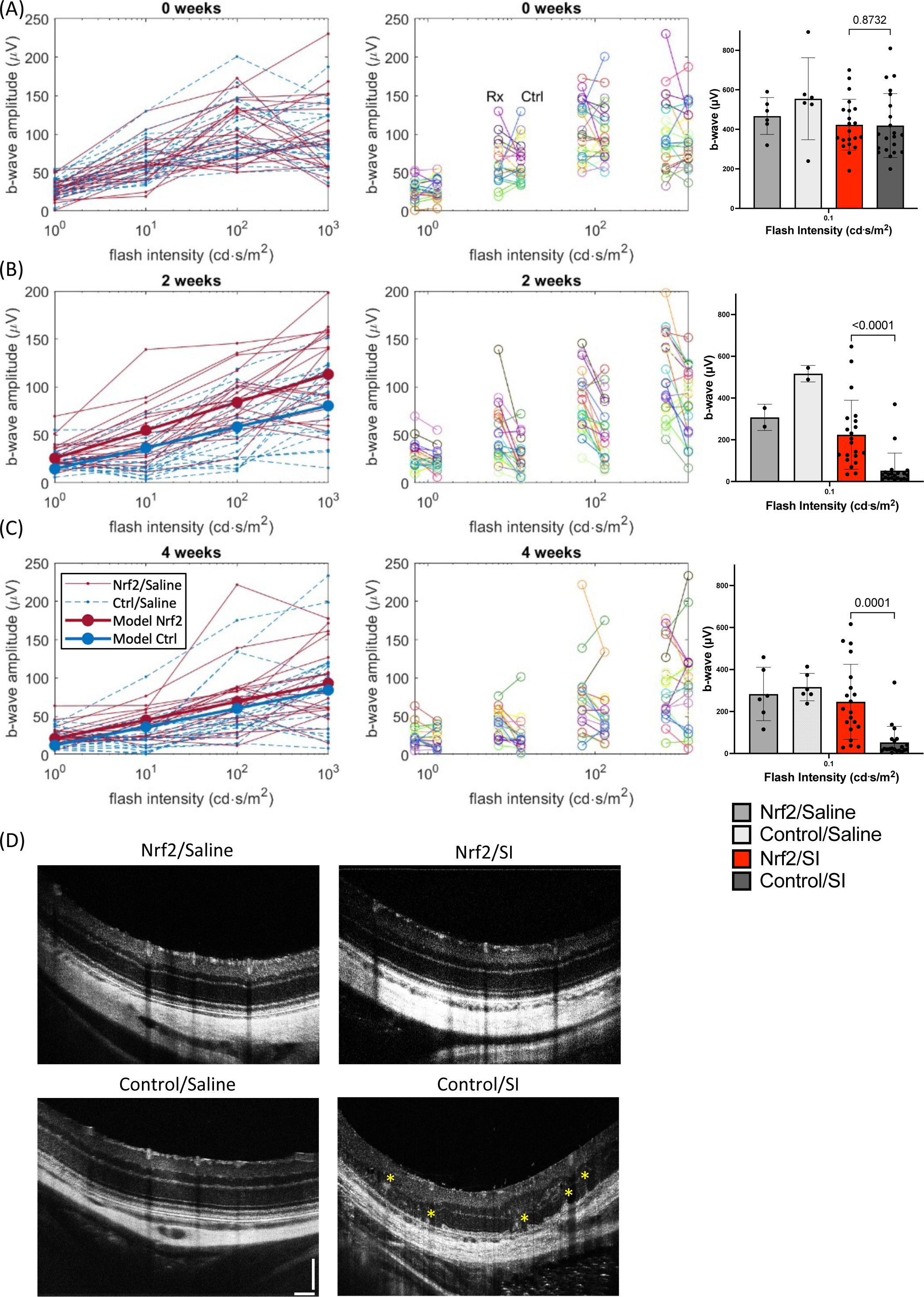
Assessment of AAV8/Best1-Nrf2 on retinal structure and visual function in rats. Rats were the same as those described in Figure 2, panels B-C. ERG b-wave amplitudes were collected at 5 different light intensities and plotted at three timepoints: baseline (before IP injection), 2 weeks post IP injection, and 4 weeks post IP injection of saline or S.I. For all photopic ERG plots, red lines denote Nrf2/SI treated eye data and blue lines denote control/SI treated eye data. For the 2 week and 4 week post IP data (panels B-C), linear mixed-effects models were fit to the data using maximum likelihood with MATLAB’s ‘fitlme’ function and the model fit curves were plotted on top of the data as thicker red/blue lines (denoted Model Nrf2 and Model Ctrl in the legend). (A) Left plot: photopic ERG data collected before IP injection with S.I. (n=22 rats). Each line represents one eye of a given rat, connected across multiple flash intensities. Middle plot: Same data as in left panel, except lines were drawn to connect the two eyes of a given rat, one of which received the Best1-Nrf2 construct while the other received a control AAV. Right plot: scotopic ERG data collected before IP injection with saline (n=6 rats) or S.I. (n=22 rats, mean ± SD, p-value is n.s., paired t-test). (B) Left/middle plots: same as panel A, except photopic ERG data was collected 2 weeks post IP injection with S.I. (n=22 rats). Right: scotopic ERG data collected 2 weeks post IP injection with saline (n=2 rats) or S.I. (n=22 rats, mean ± SD, p<0.0001, paired t-test). (C) Left/middle plots: same as panel A, except photopic ERG data was collected 4 weeks post IP injection with S.I. (n=22 rats). Right: scotopic ERG data collected 4 weeks post IP injection with saline (n=6 rats) or S.I. (n=22 rats, mean ± SD, p=0.0001, paired t-test). (D) OCT imaging collected ∼4 weeks post IP injection. Rosettes are highlighted with yellow asterisks (scale bar is 100 microns).

ERG and OCT assays were also conducted on two cohorts of mice (Figure S5-S6). Upon S.I. injection, eyes injected with control virus lost visual function, but eyes injected with AAV8/Best1-Nrf2 retained it, at least for scotopic vision in one mouse cohort (Figure S5A). In a different mouse cohort, scotopic and photopic vision did not appear to differ significantly between the AAV8/Best1-Nrf2 and control AAV-injected eyes (Figure S5B). OCT imaging showed preservation of structure (Figure S6), regardless of whether a control AAV plus GFP tracer AAV was injected in the contralateral eye (Figure S6A-B), or only a GFP tracer AAV was injected in the contralateral eye (Figure S6C). In cases where there was only partial transduction, OCT imaging showed preservation of structure only in the transduced (GFP+) region (Figure S6B).

### Injection of AAV8/Best1-Nrf2 in Adults

Dry AMD is most common among individuals over age 50.^1^ Delivery of AAV8/Best1-Nrf2 to adults was thus tested to determine if it could prevent oxidative damage when delivered to mature animals. Injecting S.I. into rodents as adults, followed by AAV8/Best1-Nrf2 subretinal injection in adults, was considered for this purpose. However, as S.I. injection kills RPE cells as early as 1-3 days post administration,^11^ and expression of AAV-encoded genes can take up to several weeks to reach maximal levels,^48,49^ this order of events was unlikely to provide a meaningful test. Instead, a regimen involving subretinal injection of AAV8/Best1-Nrf2 or control AAV into adult eyes, followed by S.I. or saline IP injection, was adopted. Subretinal injection in adult eyes does not always result in full transduction, as the inoculum does not fill the subretinal space due to a barrier imposed by RPE-outer segment interactions. This can limit the interpretation of some assays, such as ERG. However, it also provides an opportunity to analyze the effects on areas with vector transduction versus those areas without, in the same eye. By co-injecting a GFP virus, the transduced area could be identified for these analyses.

Adult C57BL/6J mice were subretinally injected with AAV8/Best1-Nrf2 + AAV8/RedO-H2B-GFP in one eye, with no injection in the contralateral eye. Alternatively, mice were injected with AAV8/Best1-6xSTOP control vector + AAV8/RedO-H2B-GFP in one eye, with no injection in the contralateral eye. IP injections with S.I. or saline were performed 1-2 weeks post vector injection. RPE flatmounts demonstrated a marked prevention of RPE cell loss in the AAV8/Best1-Nrf2 transduced area, whereas RPE flatmounts from animals injected with control AAVs showed no protection (Figure 4A). Analogously, retinal flatmounts demonstrated increased cone survival in NRF2-transduced retinal patches relative to control virus-transduced retinal patches (Figure 4B). Separate animals were harvested for RPE/retinal flatmounts for intra-eye, rather than inter-eye, RPE and cone count comparisons (Figure 5A-C). NRF2-transduced areas, identified by AAV8/RedO-H2B-GFP labeling of RPE, retained many more cells than adjacent untransduced areas in the same flatmount. This rescue effect was not seen in control AAV-transduced eyes (Figure 5A), where the transduced region was deduced from GFP+ patches in surviving RPE cells near the most peripheral edges of the flatmount, where S.I. was previously noted to show less toxicity to RPE cells. Similarly, prevention of cone loss was higher in NRF2 transduced patches of retinal flatmounts and lower in untransduced areas of the same flatmount (Figure 5B). In control virus injected eyes, transduced areas showed slightly greater cone counts than untransduced areas of the same flatmounts, but this effect was not statistically significant (Figure 5B). In eyes injected with AAV8/Best1-Nrf2 and the AAV8/RedO-H2B-GFP tracer virus, sharp boundaries delineating RPE survival were seen and were well correlated with transduction (Figure 5C). Higher magnification of areas with or without AAV transduction in the same flatmount also helped to reveal the local effects of NRF2 on RPE survival following delivery of AAV8/Best1-Nrf2 to adults.

**Figure 4:**
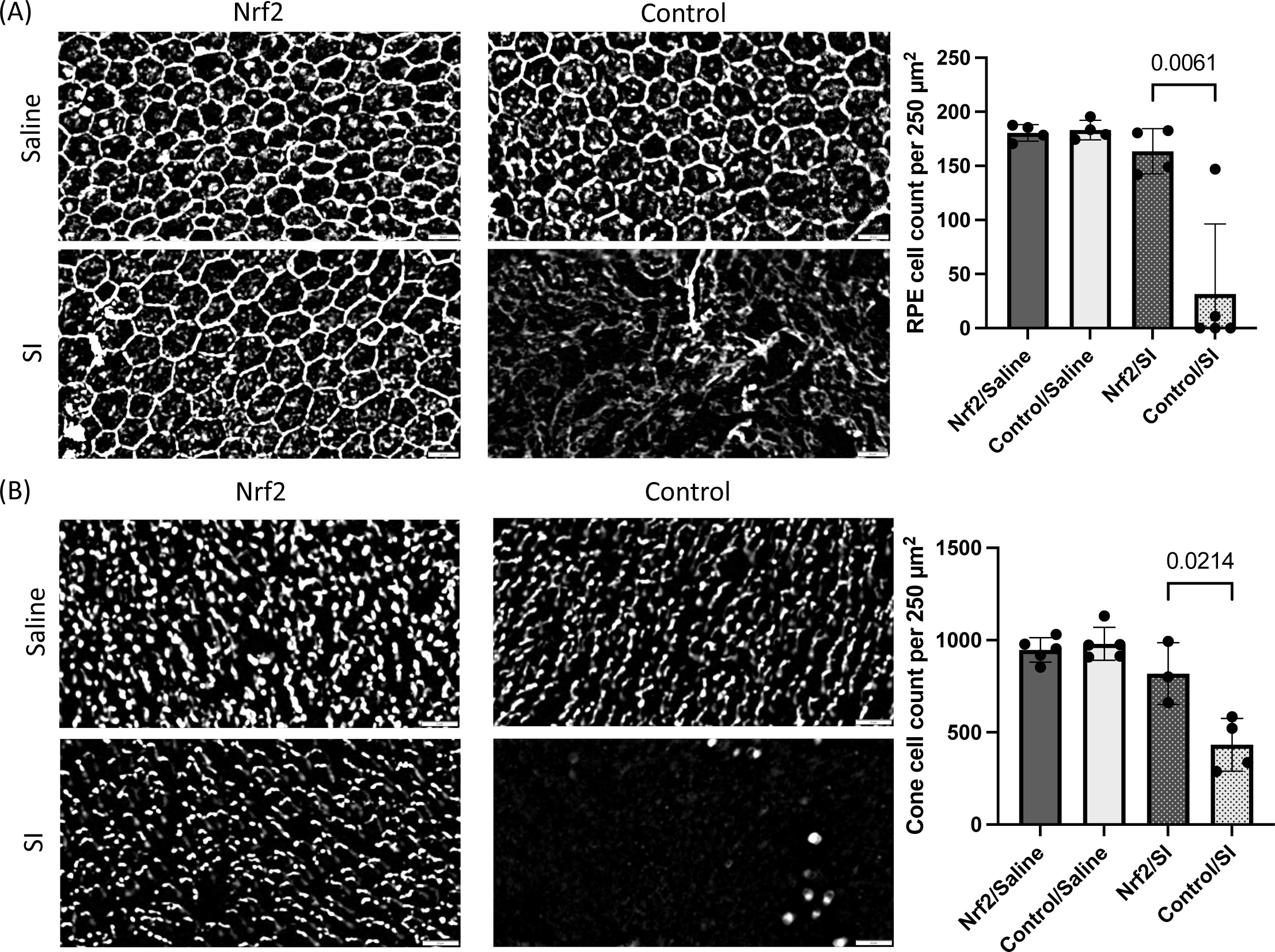
Assessment of AAV8/Best1-Nrf2 on RPE and retinal histology in mice subretinally injected as adults. C57BL/6J mice were subretinally injected at 6-13 weeks of age with AAV8/Best1-Nrf2 + AAV8/RedO-H2B-GFP in one eye (no injection in the contralateral eye), or alternatively, AAV8/Best1-6xSTOP control vector + AAV8/RedO-H2B-GFP in one eye (no injection in the contralateral eye), and IP injected with saline or S.I. at 1-2 weeks post subretinal injection. Tissues were harvested/quantified at 4-5 weeks post S.I. challenge. (A) Representative RPE flatmounts (central region) stained with phalloidin (white). Scale bar is 20 microns. Quantification of the number of RPE cells remaining is shown on the right (n=4-5 mice per bar on graph, mean ± SD, p=0.0061, unpaired t-test). (B) Representative retinal flatmounts (central region) stained with an antibody to the cone marker, CAR (white). Scale bar is 20 microns. Quantification of the number of cones remaining (see Methods) is shown on the right (n=3-5 mice per bar on graph, mean ± SD, p=0.0214, unpaired t-test).

**Figure 5:**
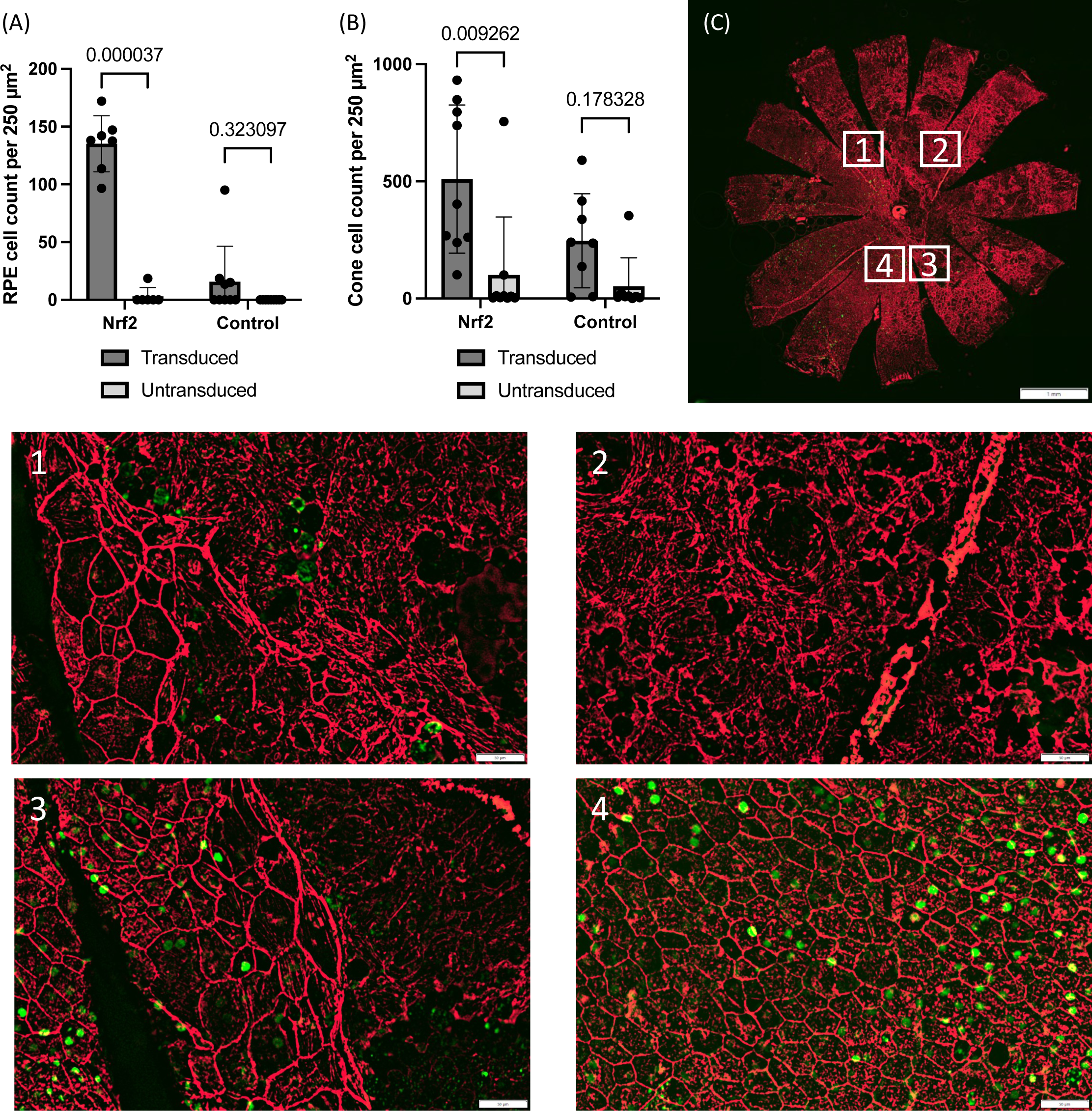
Assessment of local RPE and cone preservation in mice subretinally injected as adults. Adult mice were injected as described in Figure 4. Tissues were harvested/quantified at 11-12 weeks post IP injection of saline or S.I. (A) Quantification of the number of RPE cells remaining in the transduced or untransduced area of each RPE flatmount (n=7 NRF2/S.I. eyes and n=9 Control/S.I. eyes, mean ± SD, p<0.0001 for Nrf2 comparison, all other p-values are n.s., multiple paired t-tests with Bonferroni-Dunn multiple comparisons test). (B) Quantification of the number of cones remaining in the transduced or untransduced area of each retinal flatmount (n=9 NRF2/S.I. eyes and n=8 Control/S.I. eyes, mean ± SD, p=0.009262 for Nrf2 comparison, all other p-values are n.s., multiple paired t-tests with Bonferroni-Dunn multiple comparisons test). (C) Representative RPE flatmount from an adult mouse subretinally injected with AAV8/Best1-Nrf2 + AAV8/RedO-H2B-GFP and IP injected with S.I. at 1-2 weeks post subretinal injection is shown (scale bar is 1 mm). Phalloidin staining is in red. Within the flatmount, there is an area of live RPE cells (4), areas of transition between live transduced and dead untransduced RPE cells (1, 3), or completely absent RPE cells (2). Area of exposure to the test article is evidenced by GFP signal. Scale bars of subpanel images are all 50 microns.

ERG assays were also performed on adult mice to determine if visual function was preserved. There was no statistically significant improvement in visual function in eyes injected with AAV8/Best1-Nrf2 or control AAV relative to contralateral uninjected control eyes (Figure S7A-B). Transduction is usually 30-50% of the retina when subretinal injections are performed in adult animals. This likely leads to protection of an insufficient number of cells to be detectable by ERG, which is a relatively low-sensitivity assay.

### NRF2 pathway activation

To determine if overexpression of NRF2 resulted in activation of its canonical target genes, RNA assays were conducted. One set of neonatal mice was injected with AAV8/Best1-Nrf2 or AAV8/Best1-6xSTOP-mutGFP control vector in separate eyes. Another set was injected with AAV8/Best1-6xSTOP-mutGFP control vector vs. PBS as a vehicle control. RPE RNA was extracted from dissected eyes and quantitative PCR (qPCR) was performed to assess expression levels of NRF2, plus four canonical NRF2 target genes: NQO1, GCLC, TXNRD1, and HMOX1 (Figure 6A).^50^ Increased levels of NRF2 RNA and that of each target gene was seen in NRF2-transduced RPE relative to control-injected RPE. In addition to analysis of Nrf2 target RNAs in eyes injected with AAV8/Best1-Nrf2 and AAV8/Best1-6xSTOP-mutGFP, an analysis was done on these target RNAs in control virus injected eyes relative to vehicle injected eyes (Figure 6B). Surprisingly, one NRF2 target gene, HMOX1, was significantly downregulated in control injected RPE relative to vehicle injected RPE (Figure 6B). The other target genes, and NRF2 itself, were not significantly upregulated or downregulated relative to vehicle control RPEs in these controls (Figure 6B).

**Figure 6:**
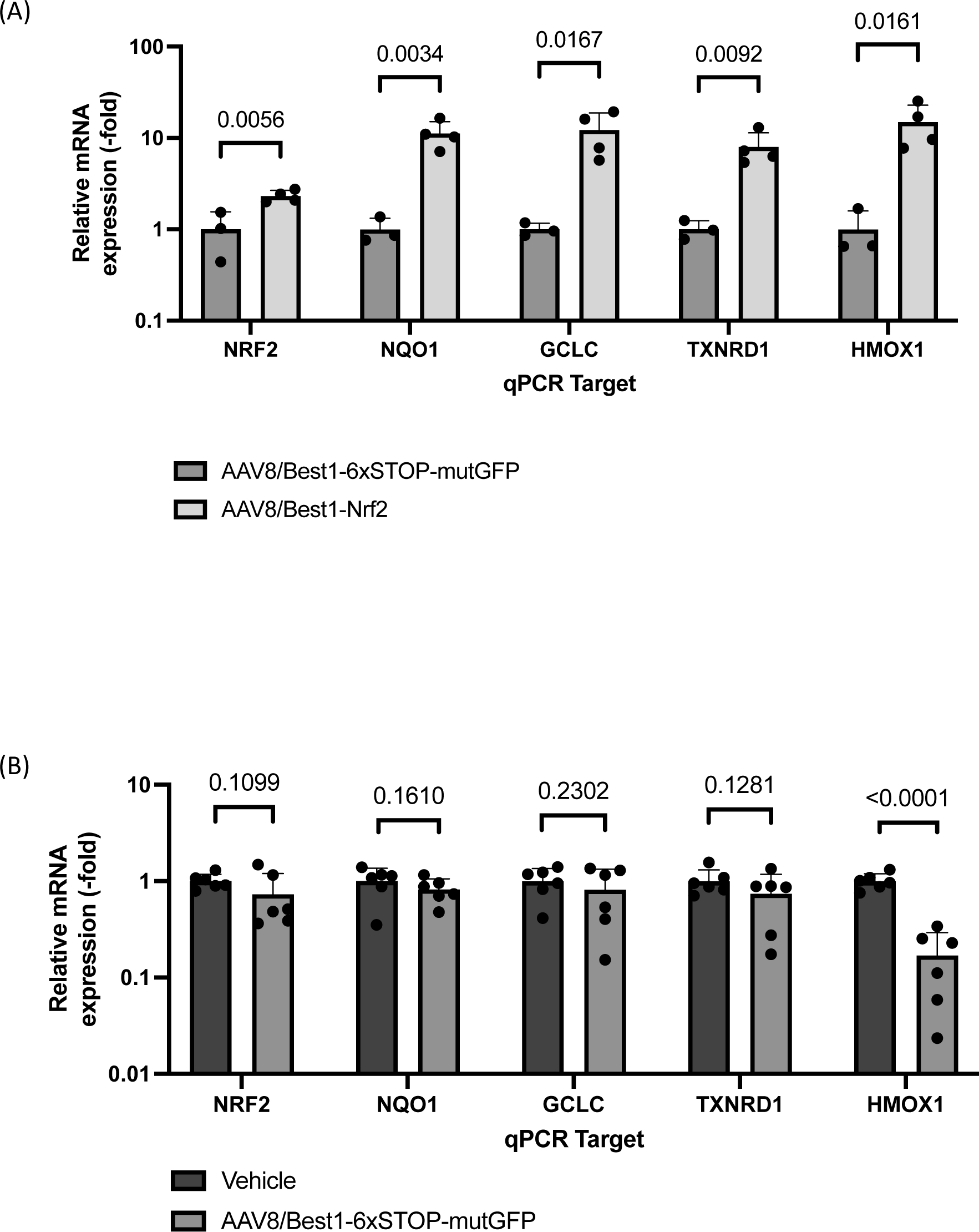
Assessment of NRF2 target gene activation. Mice were injected subretinally at birth and harvested at 7-10 weeks of age for RPE RNA extraction and qPCR quantification of NRF2 transcripts or representative NRF2 target gene transcripts. (A) RPE RNA quantification from mice injected with AAV8/Best1-Nrf2 + AAV8/RedO-H2B-GFP in one eye or AAV8/Best1-6xSTOP-mutGFP control vector + AAV8/RedO-H2B-GFP in the other eye (n=3-4 mice per target gene assessed, mean ± SD). P-values for t-tests (unpaired, equal variances, 1 tail) are shown. A sign test for the entire group was also run to give a p-value of 0.031 for the group. (B) RPE RNA quantification from mice injected with AAV8/Best1-6xSTOP-mutGFP control vector in one eye or PBS vehicle in the other eye (n=6 mice per target gene assessed, mean ± SD). P-values for t-tests (unpaired, equal variances, 1 tail) are shown.

### Long-term tolerability to AAV8/Best1-Nrf2 in rats

In addition to testing the efficacy of AAV8/Best1-Nrf2 in preserving retinal structure and function when administered to rats or mice prior to S.I. injection, the safety of this potential gene therapy was assessed long-term following injection into otherwise healthy eyes.

Two cohorts of rats were injected neonatally with virus, either at a 4e8 vg or 2e9 vg dose per eye. AAV8/Best1-Nrf2 was injected in one eye and AAV8/Best1-6xSTOPmutGFP control vector was injected in the contralateral eye. These rats were then held for ∼1 year. OCT measurements were performed on these animals at 9-10 months of age. No AAV-related toxicity was detectable, but slight, localized disruptions of retinal structure were sometimes present, likely due to injection-related trauma (Figure 7A-B). In general, OCTs revealed thinner retinas relative to images from younger rats, regardless of the treatment and dose. This is compatible with reports indicating that Sprague-Dawley rats have an approximately 50% reduction in the ONL by one year of age.^51^ ERG measurements were also performed on these same rats at 10-11 months of age. ERG results did not show significant differences between scotopic or photopic responses in animals injected with all doses and constructs (Figure 7C-D). To determine if AAV-injected eyes were comparable to non AAV-injected healthy eyes at a younger age, another cohort of rats was injected similar to the above (4e8 or 2e9 vg AAV8/Best1-Nrf2 in one eye), but the contralateral eyes were injected with PBS as a vehicle control. ERG measurements at 1-2 months showed no differences in AAV8/Best1-Nrf2 injected eyes relative to vehicle controls (Figure 7E-F).

**Figure 7:**
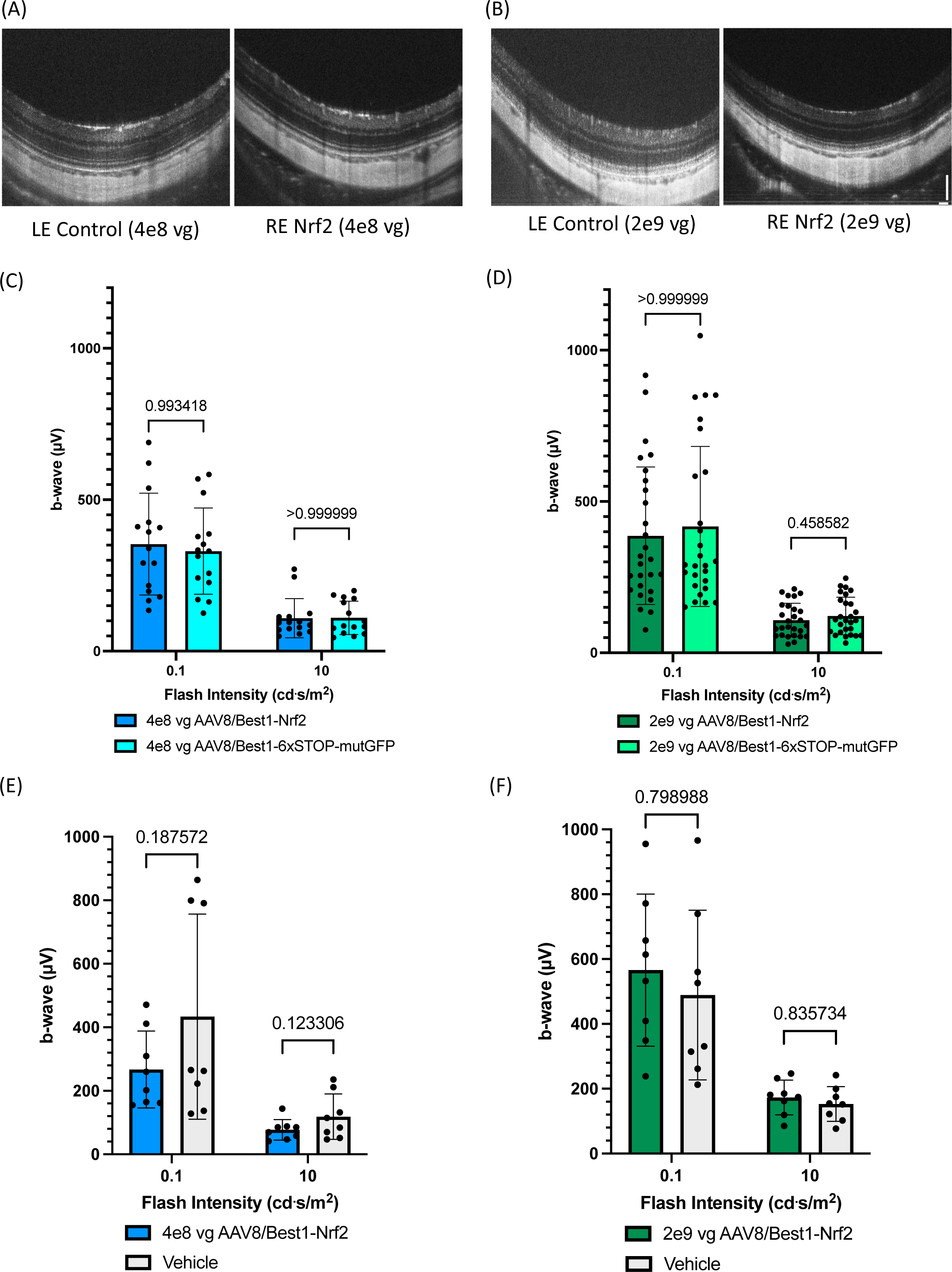
Long-term tolerability assessments for AAV8/Best1-Nrf2 in rats. OCTs were collected at 9-10 months of age and representative images are shown. ERGs were collected at 10-11 months of age and b-wave amplitudes are shown. “Scotopic” ERG used a flash stimulus of 0.1 cd.s/m^2^, while “photopic” ERG used a flash stimulus of 10 cd.s/m^2^. (A) Representative OCT images. Rats were injected with 4e8 vg of AAV8/Best1-Nrf2 or AAV8/Best1-6xSTOP-mutGFP in contralateral eyes (n=12 rats, scale bar is 100 microns). (B) Representative OCT images. Rats were injected with 2e9 vg of AAV8/Best1-Nrf2 or AAV8/Best1-6xSTOP-mutGFP in contralateral eyes (n=25 rats, scale bar is 100 microns). (C) Scotopic and photopic ERG b-wave amplitudes. Rats were injected as described in panel A (n=15 rats, mean ± SD, all p values are n.s., multiple paired t-tests with Bonferroni-Dunn multiple comparisons correction). (D) Scotopic and photopic ERG b-wave amplitudes. Rats were injected as described in panel B (n=27 rats, mean ± SD, all p values are n.s., multiple paired t-tests with Bonferroni-Dunn multiple comparisons correction). (E) Scotopic and photopic ERG b-wave amplitudes. Rats were injected with 4e8 vg AAV8/Best1-Nrf2 in one eye or PBS vehicle control in the other eye (n=8 rats, mean ± SD, all p values are n.s., multiple paired t-tests with Bonferroni-Dunn multiple comparisons correction). (F) Scotopic and photopic ERG b-wave amplitudes. Rats were injected with 2e9 vg AAV8/Best1-Nrf2 in one eye or PBS vehicle control in the other eye (n=8 rats, mean ± SD, all p values are n.s., multiple paired t-tests with Bonferroni-Dunn multiple comparisons correction).

Effects of AAV8/Best1-Nrf2 were also assessed in mice at two doses of AAV8/Best1-Nrf2, 8e7 vg/eye and 4e8 vg/eye, as shown in Figure S8. ERG measurements demonstrated no differences in scotopic and photopic conditions between AAV-injected eyes and vehicle-injected control eyes at 1-2 months of age (Figure S8A-B). Throughout the study, we used two different C57BL/6 strains: C57BL/6J, abbreviated B6J here, or C57BL/6NCrl, abbreviated B6N here, from Jackson labs or Charles River labs, respectively. This could represent a potential concern as B6N mice have been found to harbor the rd8 mutation, which causes retinal degeneration.^52^ B6J mice were used for adult subretinal injection experiments, the initial AAV8/Best1-6xSTOP-mutGFP control vector characterization, and the establishment of the 75 mg/kg S.I. IP injection model (Figures 4, 5, S1, and S7), and B6N mice were used for all other experiments requiring neonatal mouse injections of AAV8/Best1-Nrf2 (Figures 6, S3, S5, S6, and S8). To investigate if strain differences could impact the results, a comparison of histological and functional assays were carried out. RPE and retinal flatmounts, eyecup sections, and ERG were collected from 16-18 week old B6J and B6N mice (Figure S9). All histological metrics were comparable between the strains (Figure S9A-D). Quantification revealed no statistically significant difference in cone counts or ONL thickness between the strains (Figure S9B-D), although there was a slightly (statistically significant) higher number of RPE cells per 250 µm^2^ region in B6N relative to B6J mice (Figure S9A/D). ERG results between the two strains were also comparable across 5 light intensities tested (Figure S9E), indicating that for all quantitative readouts and timepoints employed in this work, the B6J and B6N strains were equivalent.

## Discussion

Oxidative stress underlies many age-related diseases,^53^ and the eye is especially susceptible to oxidative damage, due to the high oxygen demands of photoreceptors and the high light environment of the eye.^17,19^ Our lab previously demonstrated the efficacy of NRF2-expressing AAVs in reducing oxidative stress and promoting RPE and cone survival in 3 mouse models of RP.^20,21^ Hypothesizing that NRF2 might also be effective in the treatment of another blinding disease with oxidative stress, we tested it in both rat and mouse models of S.I.-induced oxidative damage. There was striking preservation of RPE and photoreceptors in AAV8/Best1-Nrf2 transduced eyes of rats and mice post S.I. challenge. Efficacy was visualized and quantified using RPE and retinal flatmounts, as well as histology of sections. Using ERG and OCT measurements over time, visual function of rods and cones, as well as retinal morphology, were seen to be retained in NRF2-treated eyes in rats. In mice, ERG assays revealed that rod function alone was significantly preserved (in one mouse cohort) or neither rod nor cone function was preserved (in another mouse cohort). These disparate outcomes in rat versus mouse cohort ERGs could be due to inherent variability from several sources: the ERG readout itself, the total number of animals assayed per cohort, the subretinal injection quality across individual animals in each cohort, and/or inherent variability in visual retention across albino vs. pigmented rodent species (discussed more below in the context of OCT). The ERG data in rats lines up with the retention of cones, as assayed by quantification of CAR-positive cells, as well as by maintenance of ONL thickness, which reflects the survival of rods.

When animals were injected with S.I. and the ONL thickness measured by OCT, ONL thinning was more obvious in mouse compared to rat, even though the injected dose of S.I. was the same (75 mg/kg). This might reflect an underlying greater sensitivity to S.I. injection in mice compared to rats, and/or it could be due to the fact that the mouse strains used (C57BL/6J or C57BL/6N) are pigmented strains compared to the albino rat strain used (Sprague-Dawley).^36^ Pigmented strains are potentially more susceptible to S.I. induced damage than albino strains, due to a reaction of melanin pigment in RPE with S.I.^54^ It could also be that C57BL/6 mouse strains are in-bred, and Sprague-Dawley rats are outbred. Genetic factors may lead to a more consistent, more severe response to S.I. injection in mice, relative to a potentially more variable, less severe effect in outbred rat strains.

Subretinal injections at birth were performed rather than adult injections in the first set of experiments due to several factors: (1) neonatal injections produce nearly full RPE and cone transduction as the inoculum readily spreads in neonates, (2) the eye can better heal after neonatal injections relative to adult injections, and (3) the expression of transgenes by AAV vector genomes is maximized after ∼4 weeks of expression, giving the vector time for NRF2 expression prior to injecting S.I. in adulthood.^48^ However, the delivery of a gene therapy agent to neonatal animals does not necessarily predict the efficacy of delivery to adult humans with ongoing disease. We therefore also tested the delivery of AAV8/Best1-Nrf2 to adult mice. We found that delivery to adults prevented S.I.-induced damage. Subretinal AAV delivery in adults, which gives localized transduction, provided an opportunity to evaluate effects of NRF2 within the same animal. Rescue was seen to be local, as RPE and cone survival was seen in transduced regions, but not in untransduced regions, of the same eye. AAV8/Best1-Nrf2 would be expected to exert cell-autonomous rescue in RPE cells, since NRF2 is a transcription factor, but it was also possible that healthier, transduced RPE cells could offer some protection to nearby untransduced RPE cells. It was difficult to precisely evaluate this possibility, but if there was non-autonomous rescue of RPE cells, it was restricted to very close neighbors of transduced cells. The protection of photoreceptors was, however, non-autonomous as they do not express genes from the BEST1 promoter. Photoreceptors might have benefitted from the better health of NRF2-overexpressing RPE cells, as RPE cells provide critical support to them. Consistent with this possibility, eyes with localized areas of transduction showed that photoreceptors benefit only in areas of RPE transduction. In addition to healthier RPE cells being better able to provide support to photoreceptors, NRF2 may lead to the upregulation and potentially the secretion of novel factors that provide local benefits for photoreceptors. Additional research is required to test this possibility.

We marked transduced areas of RPE and retinal flatmounts in nearly all experiments in the study by co-injecting the AAV8/RedO-H2B-GFP vector with AAV8/Best1-Nrf2 or a control AAV into each eye. Despite harboring the human RedO promoter, which is cone-specific,^40,55^ this AAV effectively labeled the transduced area in not just the retina, but also the RPE. RPE labeling was not due to lack of specificity of the RedO promoter, as injection of AAV8/RedO-H2B-GFP alone did not label the RPE.^55^ However, RPE labeling was detected when AAV8/RedO-H2B-GFP was co-injected with an AAV that had a promoter that was active in the RPE. We presume that concatenation or recombination of the two genomes leads to the expression of the H2B-GFP transgene in the RPE. This phenomenon has been reported by Duan et al. (2000) and Coughlin et al. (2025), with a similar interpretation regarding transcriptional cross-talk due to concatenation of co-infecting genomes.^56,57^ Interestingly, the recombination/concatenation did not occur in photoreceptors, as when an RPE promoter driving GFP and a vector with a photoreceptor promoter were co-injected, little or no GFP was seen in the photoreceptors. This may be due to the lower copy number of vectors in the photoreceptors, as when in situ hybridization was used to quantify vector genomes, there was >10 fold fewer genomes in photoreceptors relative to the copy number in the RPE, at a dose comparable to that used here.^58^

We verified the expected biological activity of NRF2 as a transcription factor by quantification of RNAs of NRF2 target genes. These results were in keeping with our previous study using RNASeq to quantify the effects of AAV8/Best1-NRF2 on RPE gene expression.^21^ We also used IHC to look for NRF2 protein. We could detect it in AAV8/Best1-Nrf2 transduced HEK293T cells (which unexpectedly express from the BEST1 promoter), but not in RPE flatmounts from AAV8/Best1-Nrf2-injected animals. Similar lack of detection was seen in a previous study of NRF2 delivered to the eye using AAV.^59^ This might be an ideal situation, as high overexpression of a powerful transcription factor could be deleterious. Consistent with this observation was the lack of toxicity in treated rats nearly one year post subretinal injection. Apparently, the BEST1 promoter resulted in enough NRF2 to drive target gene expression to protective levels, without noticeable toxicity. Given that the BEST1 promoter and NRF2 cDNA used here were derived from humans, similarly beneficial levels might be predicted if applied clinically. Our study of long-term effects of AAV control vectors with no transgene, or encoding NRF2, also provided data regarding the safety of AAV itself. AAV can cause toxicity, including within the eye, as we have documented for AAV vectors that express in the RPE in mice.^40^ It was thus particularly reassuring that no toxicity phenotypes were noted here even after nearly one year.

Regarding clinical application, we would anticipate local effects of AAV8/Best1-Nrf2 in the treatment of disease. AMD and retinitis pigmentosa damage tend to occur in patches, which grow over time.^60^ Delivery of NRF2 to healthier areas adjacent to such patches, or early before irreversible damage occurs, might slow or prevent oxidative-stress induced loss of RPE cells, much as it prevented S.I. induced damage. Delivery of NRF2 for other ocular diseases might also be beneficial, as we and others have shown for retinal ganglion cell injury.^20,61^ Other diseases of the CNS similarly have shown some benefit from NRF2 activity.^30,62^ Given these data, and the efficacy and long-term safety results demonstrated here, delivery of NRF2 to other organ systems, where oxidative damage and inflammation occur, may also prove to be beneficial.

## Materials and Methods

### Animals

Animals were handled according to protocols approved by the Institutional Animal Care and Use Committee (IACUC) of Harvard University (IACUC mouse protocol #1695, IACUC rat protocol #3495).

C57BL/6NCrl (Strain #027) untimed or timed pregnant female mice were purchased from Charles River Laboratories to obtain litters for neonatal mouse subretinal injections. Adult C57BL/6J (Strain #000664) males and females were purchased from Jackson Laboratories or bred in-house to obtain adults needed for mouse S.I. model establishment and all adult subretinal injection experiments. Untimed pregnant Sprague-Dawley female rats were purchased from Charles River Laboratories (Strain #001) and/or bred in house. Animals were bred and maintained at Harvard Medical School (HMS) on a 12-hour alternating light/dark cycle.

### AAV vector design and preparation

AAV8/Best1-Nrf2 was prepared by Spark Therapeutics and used for all experiments performed in this work. This vector was produced by triple transfection of HEK293 cells, purified, and concentrated by cesium chloride gradient purification.^63^

Control AAVs (AAV8/Best1-6xSTOP and AAV8/Best1-6xSTOPmutGFP) as well as the tracer GFP AAV (AAV8/RedO-H2B-GFP) were prepared at HMS for all experiments performed in this work. AAV8 vectors were packaged in HEK293T cells using standard transient transfection protocols and purified via iodixanol gradient ultracentrifugation as previously described.^40^ Titers of AAV batches prepared at HMS, as well as the AAV8/Best1-Nrf2 vector stock from Spark Therapeutics, were determined by running two-fold dilutions of stocks of unknown titers on a protein gel alongside a dilution series of a stock of AAV of known titer derived by PCR of genomic sequences.

### AAV Subretinal Injections

AAV was delivered by subretinal injection into P0-P2 C57BL/6NCrl pups, P0-P2 Sprague-Dawley rat pups, or 6-13 week old C57BL/6J adult mice using hand-pulled glass needles attached to an Eppendorf FemtoJet Express Microinjector (Eppendorf 5248000261, discontinued) as previously described for pups.^64^ PBS was used to dilute AAV solutions to the correct concentrations used for injections. AAV solutions were prepared with 0.01% wt/vol Fast Green FCF dye to visualize the injection bleb. Where noted, a vehicle control consisted only of 0.01% wt/vol Fast Green FCF dye in PBS injected subretinally. See Supplemental Methods for more details on needle pulling and injection procedures.

#### Typical subretinal injection doses for AAV8 vectors, unless otherwise noted

*Abbreviations (for dosing descriptions below): vg = vector genomes; NRF2 = AAV8/Best1-Nrf2 vector; Control = AAV8/Best1-6xSTOPmutGFP vector (used for all rat/mouse pup injections) or AAV8/Best1-6xSTOP vector (used for adult mouse injections); GFP = AAV8/RedO-H2B-GFP tracer vector*

##### Mouse pups

NRF2/Control vectors were each diluted to 1.6e9 vg/μL prior to injections. GFP vector was diluted to 8e8 vg/μL prior to injections. Approximately ∼0.25 μL was injected per eye.

##### Rat pups

In most experiments, NRF2/Control/GFP vectors were each diluted to 2e9 vg/μL prior to injections. Where noted in figure legends, AAV8/RedO-H2B-GFP was used at a 100-fold lower dose, so was diluted to 2e7 vg/μL prior to injections. Approximately ∼1-2 μL AAV was injected per eye.

##### Adult mice

NRF2/Control/GFP vectors were each diluted to 2e9 vg/μL prior to injections. Approximately ∼1-2 μL AAV was injected per eye.

### Intraperitoneal (IP) injections

Rats and mice were IP injected with a freshly prepared solution of S.I. (Sigma, cat # S4007-100G) diluted in sterile 0.9% saline (Fisher Scientific, #Z1376) delivered at a dose of 75 mg/kg. A similar number of rats and adult mice in each experimental cohort were set aside for vehicle control (saline only) IP injections. B6N mice injected in the study did not have a saline control cohort. Animals injected subretinally at birth were typically aged to 6-8 weeks before performing IP injections unless otherwise specified. For mice subretinally injected as adults, 6-13 week old mice were subretinally injected with AAVs, then IP injected with saline or S.I. at 1-2 weeks post subretinal injection.

### Optical Coherence Tomography (OCT)

After inducing general anesthesia, 0.5% tropicamide eye drops were used to dilate the rodent’s eyes. Eyes were then lubricated with an ophthalmic gel to avoid corneal drying. An eye was then positioned for imaging in a Phoenix Micron IV (MICRON Image-Guided OCT2 system). The adjustable holder was rotated/moved to the capture line to view the fundus image and retinal structure. The optic nerve head (ONH) was positioned on the nasal edge of the imaged area. The scan line was placed from the ONH to the periphery, and each parameter was adjusted to capture more than three OCT images per eye.

### Measurements of RNA for NRF2 target genes

After sacrificing the mice and removing their eyeballs, the cornea, iris epithelium, and lens were gently removed and discarded. The RPE layer was separated from other tissues by digestion with Dispase (Stem Cell Technologies, #07913) for a 45 min incubation at 37°C. This was followed by scraping off the choroid, adapting the protocol introduced by Shen et al.^65^

Total RNA from the RPE layer was then extracted using a Direct-zol RNA MiniPrep with TriReagent kit (Zymo, #R2051-A) according to the manufacturer’s instructions. cDNAs were synthesized using a SuperScript III First-Strand Synthesis System kit (Life Technologies Corporation, #18080051). Quantitative PCR amplifications were performed with AzuraView GreenFast qPCR Blue Mix (Azura Genomics, #AZ-2305). The relative expression values of each gene were determined by normalization to HPRT1 expression for each sample. Primer sequences were as follows:

Mouse NQO1-forward primer: AGGATGGGAGGTACTCGAATC
Mouse NQO1-reverse primer: TGCTAGAGATGACTCGGAAGG
Mouse GCLC-forward primer: GGGGTGACGAGGTGGAGTA
Mouse GCLC-reverse primer: GTTGGGGTTTGTCCTCTCCC
Mouse TXNRD1-forward primer: GGGTCCTATGACTTCGACCTG
Mouse TXNRD1-reverse primer: AGTCGGTGTGACAAAATCCAAG
Mouse HMOX1-forward primer: AAGCCGAGAATGCTGAGTTCA
Mouse HMOX1-reverse primer: GCCGTGTAGATATGGTACAAGGA
Mouse HPRT1-forward primer: CAGTCCCAGCGTCGTGATTA
Mouse HPRT1-reverse primer: TGGCCTCCCATCTCCTTCAT
*NRF2-forward primer: GGACATGGATTTGATTGACATACTT
*NRF2-reverse primer: AGTTTTTTCTGTTTTTCCAGCTCATA

*Because the NRF2 plasmid used in this study encodes human NRF2, this pair of primers was designed to target the shared part of the mRNA sequence of mouse (isoform 1) and human (isoform 1) NRF2. This pair of primers spans exons to avoid targeting genomic DNA.

### Electroretinography (ERG)

Scotopic and photopic ERGs were collected for dark-adapted rats as previously described.^66^ All animals were dark adapted for at least 2 hours prior to the start of the experiment. Anesthesia was induced by IP injection of a ketamine/xylazine cocktail (80/10 mg/kg) diluted in sterile PBS. Pupil dilation was achieved by adding one drop of Tropicamide 1% (Bausch + Lomb) solution to each eye. To keep eyes moist during ERG acquisition, PBS drops were added to each eye after corneal and reference/ground electrodes were placed. White flashes were applied to elicit ERG responses at multiple, increasingly bright intensities (0.1 cd.s/m2 for scotopic testing or 1, 10, 100, and 1000 cd.s/m2 with a 30 cd/m2 white light background for photopic testing). After ERG acquisition, Terramycin (Zoetis) antibiotic ophthalmic ointment was applied to each eye to prevent infection and keep eyes moist until each animal fully recovered from anesthesia.

### Histology

Mice and rats were euthanized by CO_2_ overdose followed by cervical dislocation. Eyes were enucleated and then processed differently depending on the desired readout: (1) the sclera/choroid/RPE layer was collected for RPE flatmounts, (2) the neuroretina layer was collected for retinal flatmounts, or (3) the eyecup (anterior chamber and lens removed, RPE + retina intact) was collected for section-based analysis. All tissues were fixed in 4% paraformaldehyde (Thermo Scientific, Cat #28908) diluted in PBS on a rocking shaker for at least 4 hours at room temperature or overnight at 4°C. After fixation, tissues were washed 3 times in PBS. Flatmount samples were directly stained with conjugated lectins or primary antibodies diluted in freshly prepared staining buffer (1% Triton X-100, 4% normal donkey serum in PBS). Samples intended for sectioning were instead frozen for processing on a cryostat as described below.

#### RPE flatmounts

After washing, RPE flatmounts were incubated in phalloidin lectin (1:100 dilution in staining buffer). Alexa Fluor 647 Phalloidin (Invitrogen #A22287), Alexa Fluor 568 Phalloidin (Invitrogen #A12380), or Alexa Fluor 488 Phalloidin (Invitrogen #A12379) were used interchangeably. Phalloidin incubations of 4 hours at room temperature or overnight at 4°C were sufficient for mouse samples, whereas overnight (or 1-3 day) incubations at room temperature were used for rat samples. After staining, samples were washed 3 times in PBS and mounted RPE side down onto coverslips (VWR, 24×50mm No. 1.5, #48393-241) in Fluoromount G mounting solution (Southern Biotech) on Superfrost Plus slides (Fisher Scientific, #22-037-246) prior to imaging.

#### Retinal flatmounts

After washing, retinal flatmounts were incubated in cone arrestin primary antibody (Millipore Sigma #AB15282, 1:3000 for mouse samples or 1:2000 - 1:3000 for rat samples) diluted in staining buffer. Incubation times for mouse samples were the same as those specified above for RPE samples, and overnight room temperature incubations were sufficient for rat samples. After staining, samples were washed 3 times in PBS, incubated in secondary antibody (Alexa Fluor® 647 AffiniPure™ Donkey Anti-Rabbit IgG (H+L), Jackson ImmunoResearch, #711-605-152, 1:750 dilution in PBS) for 2-4 hours at room temperature, washed 3 times in PBS, and then mounted with the ONL side down onto coverslips (VWR, 24×50mm No. 1.5, #48393-241) in Fluoromount G mounting solution (Southern Biotech) on Superfrost Plus slides (Fisher Scientific, #22-037-246) prior to imaging.

#### Eyecup Sections

After washing, eyecups (i.e. retina and attached RPE/sclera) were allowed to equilibrate in 10% sucrose in PBS until they sank, then they were transferred to a solution of half Optimal Cutting Temperature (OCT) compound and half 30% sucrose for at least 30 minutes at 4°C. Eyecups were frozen in cryomolds (VWR Tissue-Tek Biopsy Cryomolds, 10 mm × 10 mm x 5 mm, #25608-922) and stored at -20°C or -80°C prior to sectioning on a Leica CM 3050S cryostat (Leica Microsystems). Sections were cut at a thickness of 20 μm onto Superfrost Plus slides (Fisher Scientific, #22-037-246) prior to staining with DAPI (Invitrogen) for 15+ minutes. Fluoromount G mounting solution was then added to slides and coverslips were placed prior to imaging, as above.

For RPE65 staining, sections were cut as described above and then samples were incubated in a staining solution containing the RPE65 antibody (Abcam #ab231782, 1:250 dilution in staining buffer) and DAPI for 5-6 hours at room temperature. Sections were then washed with PBS and incubated in secondary antibody solution (Alexa Fluor® 488 AffiniPure™ Donkey Anti-Rabbit IgG (H+L), Jackson ImmunoResearch, #711-545-152, 1:750 dilution in PBS) for 2 hours at room temperature prior to a final wash with PBS. Fluoromount G mounting solution was then added to slides and coverslips were placed prior to imaging, as above.

#### Whole Eye H&E Preparation

Eyes were harvested from rats (4-5 weeks post IP injection) and immediately placed in 4% paraformaldehyde (with a corneal slit to allow for better fixative penetration) for overnight incubation at 4°C prior to delivering samples to the Schepens Eye Research Institute of Mass Eye and Ear Morphology Core for paraffin embedding, sectioning to 6 µm, and hematoxylin & eosin staining. A separate set of eyes harvested from S.I. injected mice (12-13 weeks post S.I. injection, no corneal slit made) was fixed overnight in Davidson’s fixative at 4°C prior to delivering samples to the Schepens Eye Research Institute of Mass Eye and Ear Morphology Core for paraffin embedding, sectioning, and hematoxylin & eosin staining. Davidson’s fixative was prepared by mixing 300 mL 95% ethyl alcohol, 200 mL 10% neutral buffered formalin, 100 mL glacial acetic acid, and 300 mL distilled water. Uninjected 7-9 week-old B6J control eyes (fixed with either Davidson’s fixative or 4% paraformaldehyde overnight at 4°C) were also submitted for identical H&E processing by the Morphology Core as healthy controls for the mouse S.I. samples above.

### Image Acquisition/Analysis

All coverslipped flatmounts and sections were imaged on an Olympus VS200 Slide Scanner with a UPlan X Apo 10x/0.4 Air objective. Images were analyzed using ImageJ/Fiji software (NIH) for RPE and retinal flatmounts, or OlyVIA (Olympus Life Sciences) for eyecup section ONL thickness analysis. For RPE, 250×250 µm^2^ boxes were drawn in midperipheral areas around the flatmount, cells were manually counted in these four regions, and the median RPE count from these four regions was reported. Cones were counted within four similarly sized boxes drawn on retinal flatmounts, but an automated ImageJ/Fiji macro counted them and the median cone count from these four regions was reported per flatmount. For mouse eyecup sections, ONL thicknesses were measured ∼1 mm away from the optic nerve on either side (mid-peripheral region), and the average of the two thickness measurements was reported. For rats, thicknesses were also typically measured ∼1 mm away from the optic nerve unless that specific region was damaged (e.g., by cryosectioning), in which case the distance was extended up to ∼2 mm away from the optic nerve to find a better quality region suitable for thickness quantification (rat tissues are larger so this distance still roughly corresponds to the mid-peripheral region). See Supplemental Methods for more specific details regarding image quantification in full vs. partially transduced flatmounts.

### Statistics

All statistical analyses were performed using GraphPad Prism Version 10, Microsoft Excel, or MATLAB. MATLAB was only used for linear regression analysis of photopic ERG data. For all figures, n refers to individual eyes. All comparisons made between eyes from two different groups of animals used an unpaired Student’s t-test. Comparisons between contralateral eyes of the same animals were analyzed using a paired Student’s t-test. When one eye was damaged or otherwise unquantifiable, causing a loss of pairing, these values were ignored for paired t-test calculations. All statistical analyses were performed using a p-value cutoff of 0.05. Any p-values above this cutoff are reported as not significant or n.s. For RPE and cone counts, median cell counts were plotted as individual data points on graphs, but the error bars shown in each plot represents the mean ± SD. ERG data was analyzed using a combination of paired t-tests (for scotopic data) or regression modeling (for photopic data at multiple flash intensities, see Supplemental Methods for more specific details on how these regression analyses were performed).

## Data Availability Statement

Raw data supporting the findings of this study can be made available upon request. MATLAB code and raw data used to generate the linear mixed-effects models of photopic ERG data shown in Figure 3 and Figure S5 are available here: https://github.com/rickborn/NaIO3-Paper.git.

## Acknowledgments

We would like to thank the MicRoN microscopy core at Harvard Medical School (Dr. Paula Montero Llopis, Dr. Praju Vikas Anekal, and Dr. Adrienne Wells) for their expert microscopy advice. This study was funded by Howard Hughes Medical Institute, Spark Therapeutics, and the National Eye Institute, NIH F31 Ruth L. Kirschstein NRSA (A.G.).

## Author Contributions

A.G., S.W., C.L.C., and V.H. designed research; A.G., S.W., A.D., D.L., C.W., L.L., C.H., S.R.Z., and G.W. performed research, A.G., S.W., A.D., D.L., C.W., R.T.B., and C.L.C. analyzed the data, and A.G. and C.L.C. wrote the paper with feedback and support from R.T.B., K.K., L.B.R., and V.H. R.T.B. advised and/or performed the statistical analyses.

## Supplemental Methods

### Image Analysis

#### RPE flatmounts

RPE flatmounts were quantified as follows. A custom Fiji macro was written to extract four equally sized (∼250 µm^2^) boxes containing RPE cell areas from a full-sized RPE flatmount (phalloidin channel) image. The locations of these four boxes were dictated by four lines drawn by the investigator from the outer RPE boundary to the optic nerve head in the RPE flatmount, within the GFP+/transduced region for S.I. images or anywhere around the flatmount for saline images. The midpoints of these four lines were automatically calculated and used to define one corner of each of the four box placements. Care was taken by the investigator to ensure that these lines were drawn such that boxes would be roughly equally spaced around the flatmount and regions of out of focus imaging or tissue processing damage were avoided. The number of RPE cells remaining in each of the four regions placed by the macro was manually counted by the investigator using a built-in Cell Counter feature in Fiji. The overall median RPE count from these four regions was then calculated and reported. In few cases, a boxed region was suboptimal for manual quantification due to out of focus imaging or weak staining and was thus omitted from the median calculation.

When adult mouse injections were performed, partial transduction was the expected result, as shown by GFP. In these cases, for intra-eye injections (Figure 5), this macro was run twice on the same flatmount, once with four lines drawn from the outer edges to optic nerve head traversing a *transduced* area of the RPE flatmount, and once with four lines drawn from the outer edges to optic nerve head traversing an *untransduced* area of the RPE flatmount, using the GFP channel to define transduced areas. For control vector injected RPEs that showed RPE cell death in the entire central region of the flatmount, “transduced area” was defined by first identifying H2B-GFP+ peripheral (ciliary margin-adjacent) RPE cells which were generally not entirely killed by S.I. injection. Lines were drawn starting from RPE edges directly adjacent to those surviving peripheral GFP+ cells to the optic nerve head area in the center of the flatmount.

#### Retinal flatmounts

Cone counts for rats and mice injected with saline from the Figure 1B/E and Figure S1B/E cohorts of animals were previously published by our group to provide an estimate of cone density in wild type animals in a concurrent study.^64^ Retinal flatmounts for rats and mice in this study were quantified as previously described for rat retinas in that prior study.^64^ In cases where there was full transduction, a custom Fiji macro again extracted four equally sized (∼250 µm^2^) boxes containing labeled cones from a given full-sized retinal flatmount image. The locations of these four boxes were again placed at the midpoints of four separate lines drawn by the investigator from the outer retinal boundary to the optic nerve head in the full-sized retinal flatmount image itself. Instead of manually counting cells in each of the four boxes drawn on the flatmount, the macro automatically thresholded and counted the number of puncta in each square box and output four counts (one from each box). The overall median of these four counts was reported for cone survival assays.

When there was partial transduction, the sample was either excluded (see Sample Exclusion Criteria below), or the macro was run twice on the same flatmount, once with four lines drawn from the outer edges to optic nerve head traversing a *transduced* area of the retinal flatmount, and once with four lines drawn from the outer edges to optic nerve head traversing an *untransduced* area of the retinal flatmount (e.g., for adult mouse subretinal injections where partial transduction was the expected injection outcome). Transduced areas were marked by AAV8/RedO-H2B-GFP co-injection, which labels cones, even in control AAV/S.I. challenged eyes since cones do not entirely die after S.I. administration in the models and harvest timepoints investigated.

#### Whole Eye or Eyecup Sections

To calculate ONL thickness in whole eye H&E sections in mice, a line was drawn from the optic nerve head to the RPE-facing side of the ONL on either side (1000 µm in length). The ONL thickness in that region was measured using OlyVIA visualization software. The thickness of the ONL on both sides of the optic nerve head in the eyecup were averaged and this average was plotted. If one side of the optic nerve head was damaged from tissue processing issues, the ONL thickness on that side was not calculated and the single ONL thickness value on the other side of the section was reported as the value for that eye.

For eyecup sections in rats, a similar approach was used, except a line ranging anywhere from 1000-2000 µm in length (all within what could be considered the “mid-peripheral” region of rat sections) was drawn and a representative region of the ONL was chosen for measurement within this distance to circumvent any tissue processing/cryostat-related damage that may have been present at exactly 1000 µm away from the optic nerve head.

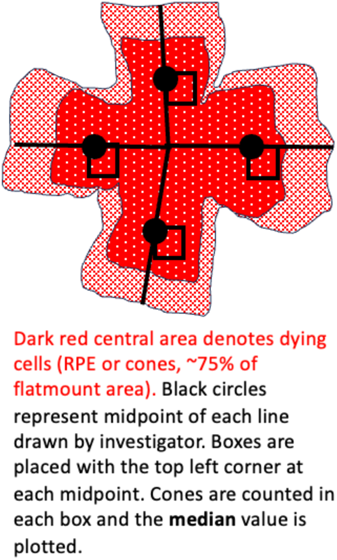

### Sample Exclusion Criteria

Flatmounts were excluded from both eyes of animals that appeared to have an IP injection issue or general lack of S.I. induced damage in its control AAV/S.I. injected eye. Some animals were missing a retina, RPE, or section quantification because the tissue in that eye was too torn or damaged post-dissection to flatmount/quantify, or there was an antibody/IHC staining or imaging issue (rare). Lastly, for rats treated with S.I. for which the NRF2 eye was not completely transduced (as determined by inspection of RedO-H2B-GFP signal in the retinal flatmount), those retinal flatmounts were excluded.

### Additional information on Subretinal Injections

Animals were administered the analgesic buprenorphine (0.05-0.1 mg/kg, Par Pharmaceutical, NDC 42023-179-01) subcutaneously by a 31G insulin syringe (BD cat #SY8290328291) just prior to the subretinal injection, and a drop of Proparacaine Hydrochloride ophthalmic solution, USP 0.5% (Akorn, NDC 17478-263-12) into each eye post subretinal injection. For adult rodent subretinal injections, a small amount of Terramycin ointment (Zoetis, NADA #8-763) was additionally added to each eye post subretinal injection as a lubricant and antibiotic.

#### Mouse/Rat Pup Injections

Pups were briefly anesthetized by cryoanesthesia on a paper towel on ice. The palpebral fissure was opened with a 30G needle. The eye was gently popped out of the socket with blunt forceps, and AAV diluted in PBS was injected into the subretinal space. Both left and right eyes were used, with the contralateral eye serving as a control for the experimental condition (e.g., left eye would receive AAV8/Best1-Nrf2, right eye would receive an equivalent dose of AAV8/Best1-6xSTOP-mutGFP or AAV8/Best1-6xSTOP control AAV). To avoid bias in injection coverage in either the left or right eyes, some animals received AAV8/Best1-Nrf2 in the left eye and the remaining received it in the right eye. Where noted, AAV8/RedO-H2B-GFP was co-injected as an injection tracer to demarcate the transduced area of flatmounts.

#### Adult mouse injections

Adult mice were given an IP administration of ketamine (80 mg/kg) and xylazine (10 mg/kg) diluted in PBS for temporary anesthesia. The eye was held open and a 30G needle was used to create a small hole at the base of the cornea (at the corneal-scleral interface) for a hand-pulled needle to enter through. AAV was then injected into the subretinal space. For each animal, the same AAV solution (AAV8/Best1-Nrf2 or AAV8/Best1-6xSTOP control AAV, along with AAV8/RedO-H2B-GFP as an injection tracer) was injected into either the right eye only, or both the left and right eyes. Right eyes were preferred for adult subretinal injections due to potentially better injection coverage in the right eye over the left eye. 1-2 weeks post subretinal injection, adult mice were administered saline or S.I. via IP injection. Eyes were harvested for histological analyses between 4-5 or 11-12 weeks post IP injection as noted in the figure legends.

*NOTE 1: The AAV8/Best1-6xSTOPmutGFP control vector was used for neonatal rat/mouse subretinal injections, whereas the AAV8/Best1-6xSTOP control vector was used for adult mouse subretinal injections. These vectors are considered interchangeable for experimental purposes because saline control animals in their respective experiments showed minimal/no RPE toxicity regardless of the control vector used. See also Figure S3 for an AAV8/Best1-6xSTOPmutGFP ONL thickness characterization, indicating its safety/neutrality as a control vector*.

*NOTE 2: For some neonatal mouse experiments, no control vector was injected, and the control eye consisted only of an AAV8/RedO-H2B-GFP injection (as noted in figure legends)*.

### Preparation of glass needles for subretinal injections

Needles for subretinal injection were made from Wiretrol II glass capillaries (Drummond #5-000-2005). Capillaries were pulled on a Model P-97 Flaming/Brown Micropipette Puller from Sutter Instrument Co. with the following settings: P (Pressure Setting) = 500, Heat = 645, Pull = 30, Vel. (Velocity) = 45, Time = 50. Needles were sharpened on a Narishige Micro Grinder (#EG-401) with the angle set to 30° and grinding disk speed to ∼70. As the eyepiece includes a micrometer, needle tips were adjusted to be ∼1 µm thick and sharpened until they had a visible bevel and point.

### Linear Mixed-effect Modeling of Photopic ERG Data

To analyze photopic ERG data shown in **Figure 3** and **Figure S5**, we used a mixed-effects linear regression model (Fitzmaurice, Laird, and Ware 2011; Laird and Ware 1982), with fixed effects for the covariates of interest and random effects for each rat, in order to account for the repeated ERG measurements for each animal across flash intensity and time, and for eye-within-rat to account for the fact that each animal served as its own control: one eye received a vector expressing the NRF2 gene and the other a control vector. In Wilkinson notation (Wilkinson and Rogers 1973), the full model was:

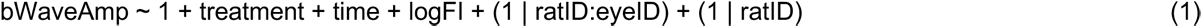

where *bWaveAmp* is the amplitude of the ERG b-wave (in µV), *treatment* is an indicator variable set to ‘1’ for eyes that received an intraocular injection of Nrf2 AAV and ‘0’ for control AAV, *time* represents the time at which the ERG recordings were made (0, 2 or 4 weeks after the IP injection of S.I.), logFI was the log10 of the flash intensity (in cd·s/m2), and the terms (1 | ratID:eyeID) and (1 | ratID) represents the random effect for eye-within-rat and rat, respectively.

The full model was fit to the data using the ‘fitlme’ function in MATLAB (MathWorks, Natick, MA), using maximum likelihood. The results are shown in **Supplemental Table 1**.

**Supplemental Table 1.** Results for full model (1). Model AIC = 4932; adjusted R^2^ = 0.52.

| Name | Estimate | Std. Error | t-Stat | DF | p Value | 95% CI (lower) | 95% CI (upper) |
| --- | --- | --- | --- | --- | --- | --- | --- |
| y-intercept | 29.57 | 4.20 | 7.03 | 500 | $6.65 \times 10^{-12}$ | 21.31 | 37.84 |
| treatment | 10.26 | 3.08 | 3.33 | 500 | $9.41 \times 10^{-4}$ | 4.20 | 16.32 |
| time | -4.81 | 0.84 | -5.70 | 500 | $2.02 \times 10^{-8}$ | -6.47 | -3.15 |
| logFI | 24.65 | 1.22 | 20.25 | 500 | $5.23 \times 10^{-67}$ | 22.25 | 27.04 |

All the model parameters were highly statistically significant (p < 0.001), and it accounted for just over 50% of the variance, after adjusting for the number of parameters in the model. The magnitude of the *treatment* parameter tells us that, on average, across all flash intensities and time-points, the Nrf2 treated eyes had a b-wave amplitude of about 10 µV greater than the control eyes, with a 95% confidence interval of [4.20, 16.32]. The negative value of the *time* parameter indicates that there was a progressive decrease in b-wave amplitude after sodium iodate treatment. We also ran models with interaction terms for ‘*treatment* x *time’* and ‘*treatment* x *logFI*, but neither of the interaction terms was significant.

Because we were not interested in the effect of time, *per se*, we also fit a reduced model to the data from each week’s recordings separately. The reduced model was the same as (1) but omitted *time* as a covariate:

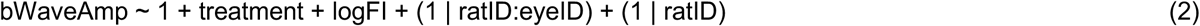

As expected, there was no detectable treatment effect at baseline (p = 0.94). The largest treatment effect was at week 2 (**Supplemental Table 2**), while the effect at 4 weeks was smaller and barely statistically significant (**Supplemental Table 3**).

**Supplemental Table 2.** Results for reduced model (2) fit to 2-week data. Model AIC = 1588; adjusted R^2^ = 0.71.

| Name | Estimate | Std. Error | t-Stat | DF | p Value | 95% CI (lower) | 95% CI (upper) |
| --- | --- | --- | --- | --- | --- | --- | --- |
| y-intercept | 9.17 | 5.21 | 1.76 | 165 | 0.080 | -1.11 | 19.45 |
| treatment | 21.95 | 4.09 | 5.36 | 165 | $2.71 \times 10^{-7}$ | 13.87 | 30.02 |
| logFI | 25.57 | 1.62 | 15.83 | 165 | $6.37 \times 10^{-35}$ | 22.38 | 28.76 |

**Supplemental Table 3.** Results for reduced model (2) fit to 4-week data. Model AIC = 1570; adjusted R^2^ = 0.61.

| Name | Estimate | Std. Error | t-Stat | DF | p Value | 95% CI (lower) | 95% CI (upper) |
| --- | --- | --- | --- | --- | --- | --- | --- |
| y-intercept | 12.37 | 6.59 | 1.88 | 157 | 0.0624 | -0.65 | 25.40 |
| treatment | 8.82 | 4.45 | 1.98 | 157 | 0.0492 | 0.03 | 17.60 |
| logFI | 24.01 | 1.99 | 12.07 | 157 | $3.82 \times 10^{-24}$ | 20.08 | 27.94 |

For the week 2 data, the slopes of the Nrf2-treated eyes appeared steeper than those of the controls (**Figure 3B)** so we fit an additional model that included an interaction term for *treatment* and *logFI*:

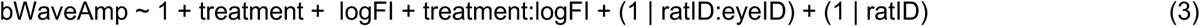

The interaction term for the 2-week data was statistically significant (7.25 µV / decade flash intensity, 95% CI [1.0, 13.5], p = 0.023; model adjusted R^2^ = 0.72), indicating that the eyes treated with Nrf2 had a steeper slope to the relationship between b-wave amplitude and flash intensity than did control eyes (Nrf2, slope = 29.2 µV / decade flash intensity; Ctrl, slope = 21.9 µV / decade flash intensity). The difference in slopes is shown by the heavy lines representing the model fits (**Figure 3B**). The interaction term was not significant for the 4-week data (p = 0.87; **Figure 3C**).

Because the treatment-vs-control data were paired within rat (i.e. for each rat that received an IP injection of the sodium iodate toxin, one eye received Nrf2 and the other eye received a null construct), we further analyzed the *differences* between eyes, which allowed us to remove sources of *shared variability* due to differences among rats in the response to the sodium iodate treatment. Such sources of shared variability might include procedural variability, such as the actual amount of toxin injected into a given rat, as well as biological variability in each rat’s weight, metabolism and RPE response to the toxin. Such shared variability is evident in **Figure 3**, particularly for the week 2 data, where the values for the 2 eyes in each rat are connected by a line. We also plotted these between-eye differences for each rat across different flash intensities (see plot at right). The plots of the 0-week data are randomly distributed about the line of y = 0, which is the null value for the paired comparisons. For the 2- and 4-week data, the fact that the preponderance of the lines are above y = 0 is indicative of the treatment effect.

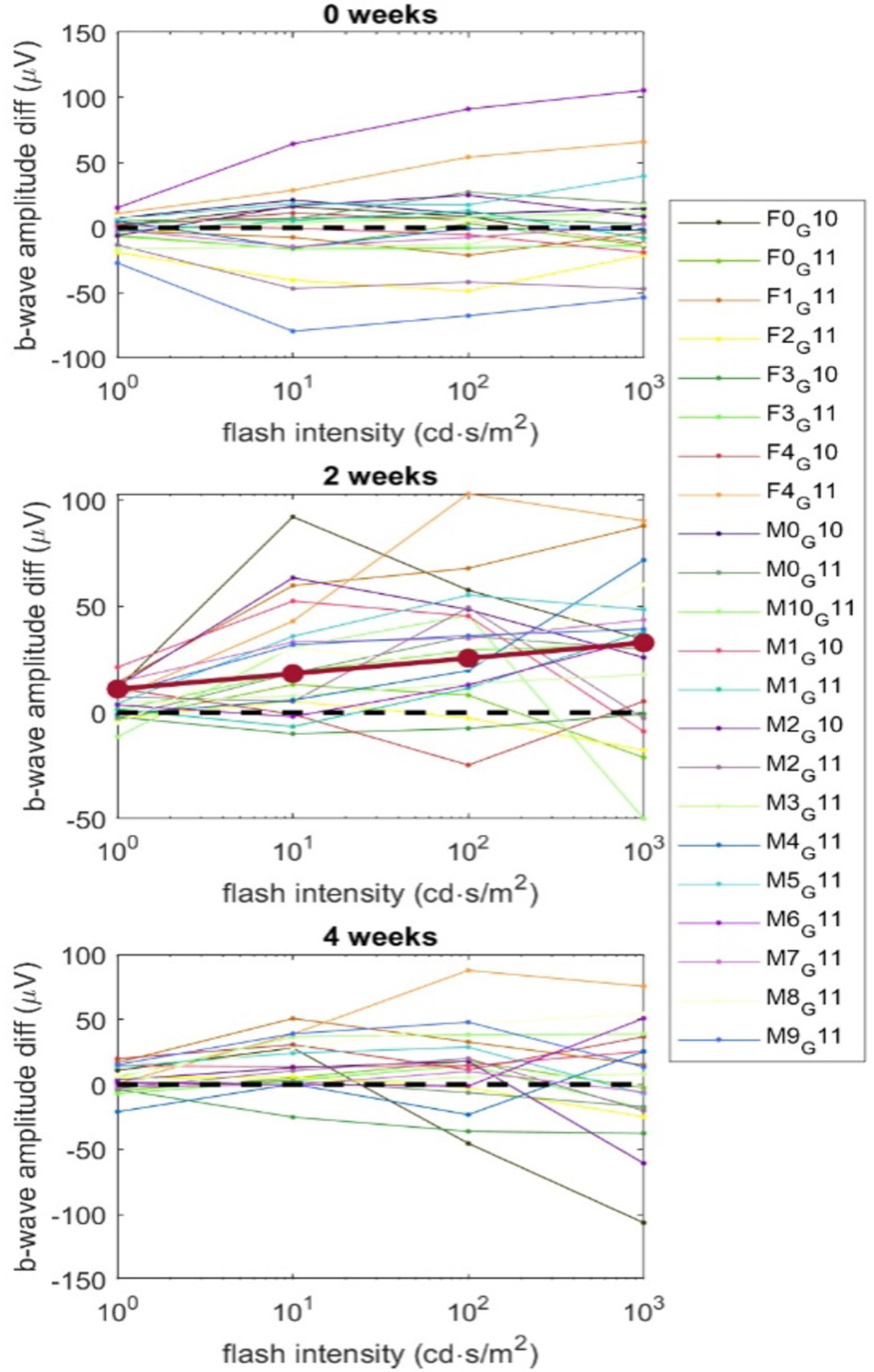

Between-eye difference plots for rats in Figure 3. We quantified this treatment effect by fitting a simplified mixed effects model to the data in the right-hand column of Figure 1:

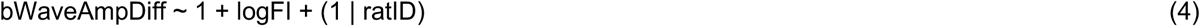

Where the dependent variable is now the within-rat difference in b-wave ERG amplitude between the treatment and control eyes. The random effect for ratID allows each rat to have its own y-intercept (as for the previous mixed effect models). The results of this fit are shown in Supplemental Table 4.

**Supplemental Table 4.**
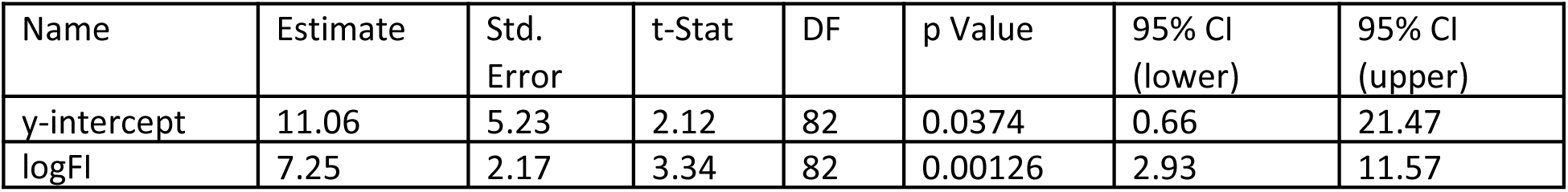
Results for reduced model (4) fit to 2-week data. Model AIC = 789; adjusted R^2^ = 0.33.

In this case, because we are operating on differences, both the y-intercept and the slope (*logFI* coefficient) are of interest: the y-intercept represents a significant offset due to the treatment, and the positive coefficient for *logFI* (slope) indicates that the effect of the treatment increases with flash intensity. In fact, the values for these two coefficients are identical to the values for the *treatment* and interaction term (*treatment* x *logFI*) in model (3). However, because the within-animal subtraction effectively removed a source of shared variability, the standard errors for the coefficients in model (4) are smaller than those of model (3), resulting in lower p-values: 0.0012 vs. 0.023, for the slope of model (4) vs. the interaction term of model (3).

Importantly, all of the above analyses are in agreement and consistent with the interpretation of a large protective effect of the Nrf2 treatment at 2 weeks and a more moderate effect at 4 weeks, both of which are highly statistically significant.

Finally, we performed two diagnostic tests of the model fits, which, taken together, indicate that our use of the linear model was appropriate. A plot of the raw residuals vs. the fitted values below (left plot) shows reasonable homoscedasticity and linearity, while the Q-Q plot displays the normality of the residuals (right plot).

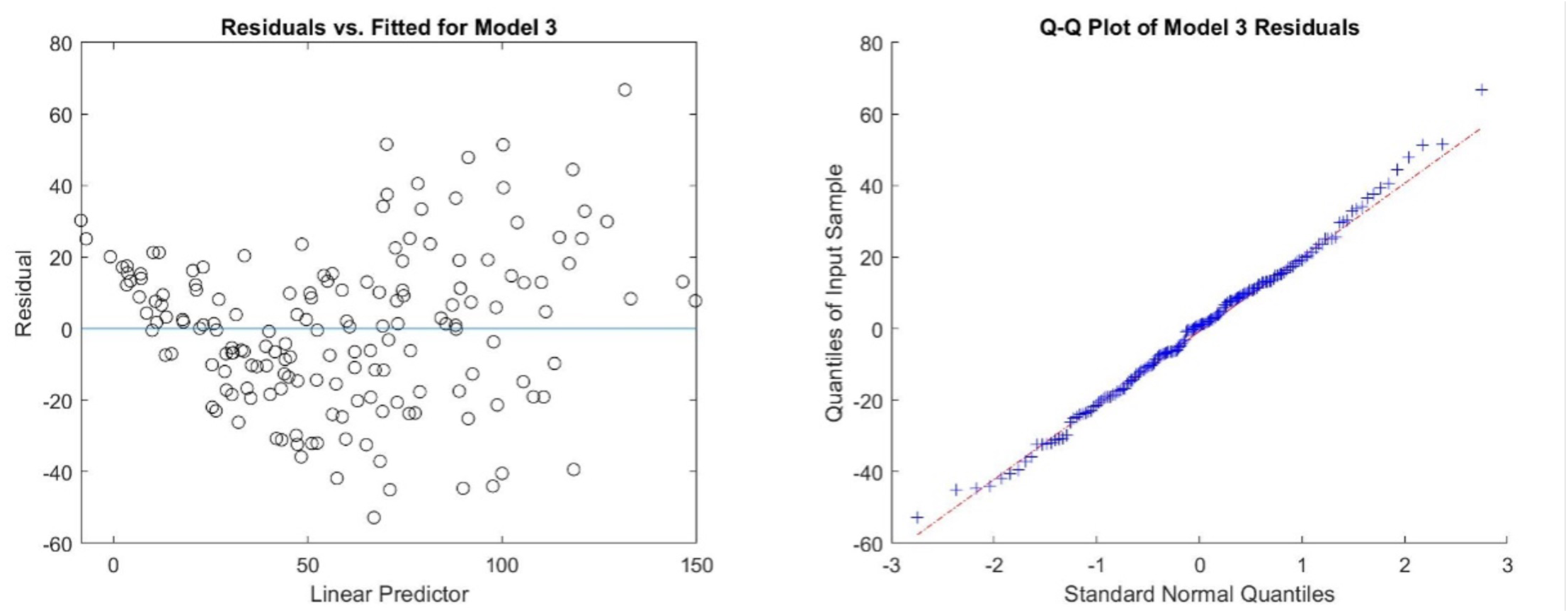

Regression diagnostics for model (3) fit to the 2-week data.

The raw data, analysis code and results are available at: https://github.com/rickborn/NaIO3-Paper.git

---

**Figure S1:**
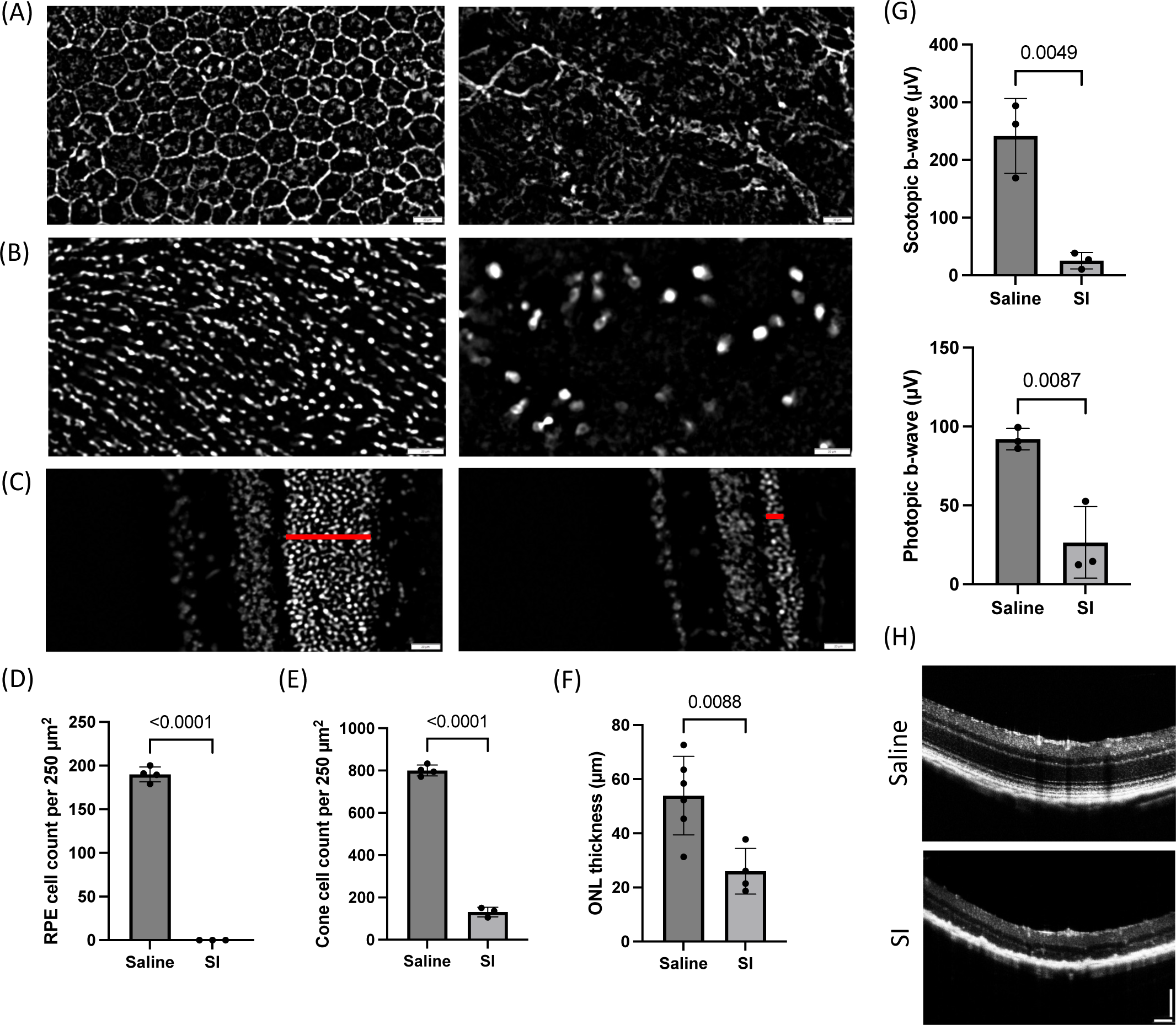
Assessment of RPE and retinal histology after IP injection of S.I. in mice. C57BL/6J mice were IP injected with saline or S.I. at ∼6-8 weeks of age. Tissue was harvested 4-7 weeks post S.I. injection. (A) Representative RPE flatmounts (central region) from a saline (left) or S.I. (right) IP-injected mouse. Flatmounts were stained with phalloidin (white). Scale bar is 20 microns. (B) Representative retinal flatmounts (central region) from a saline (left) or S.I. (right) IP-injected mouse. Flatmounts were stained with an antibody to CAR (white). Scale bar is 20 microns. (C) Representative eyecup sections (mid-peripheral region) from a saline (left) or S.I. (right) IP-injected mouse. Sections were stained with DAPI (white). Red bar indicates the ONL. Scale bar is 20 microns. (D) Quantification of the number of RPE cells (see Methods, n=3-4 eyes per group, mean ± SD, p<0.0001; unpaired T test). (E) Quantification of the number of cones (see Methods, n=3-4 eyes per group, mean ± SD, p<0.0001; unpaired T test). (F) Quantification of the average ONL thickness from eyecup sections (see Methods, n=4-6 per group, mean ± SD, p=0.0088; unpaired T test). (G) Scotopic ERG b-wave amplitudes (at a 0.1 cd.s/m^2^ flash stimulus, left) and photopic ERG b-wave amplitudes (at a 10 cd.s/m^2^ flash stimulus, right) for mice injected with saline or S.I. were collected 3-4 weeks post IP injection (n=3 mice per group, mean ± SD, p=0.0049 for scotopic and p=0.0087 for photopic, unpaired T test for each plot). (H) Representative OCT images for a saline IP-injected mouse (top panel) or an S.I. IP-injected mouse (bottom panel) at 4 weeks post IP injection (n=3 mice per group, scale bar is 100 microns).

**Figure S2.**
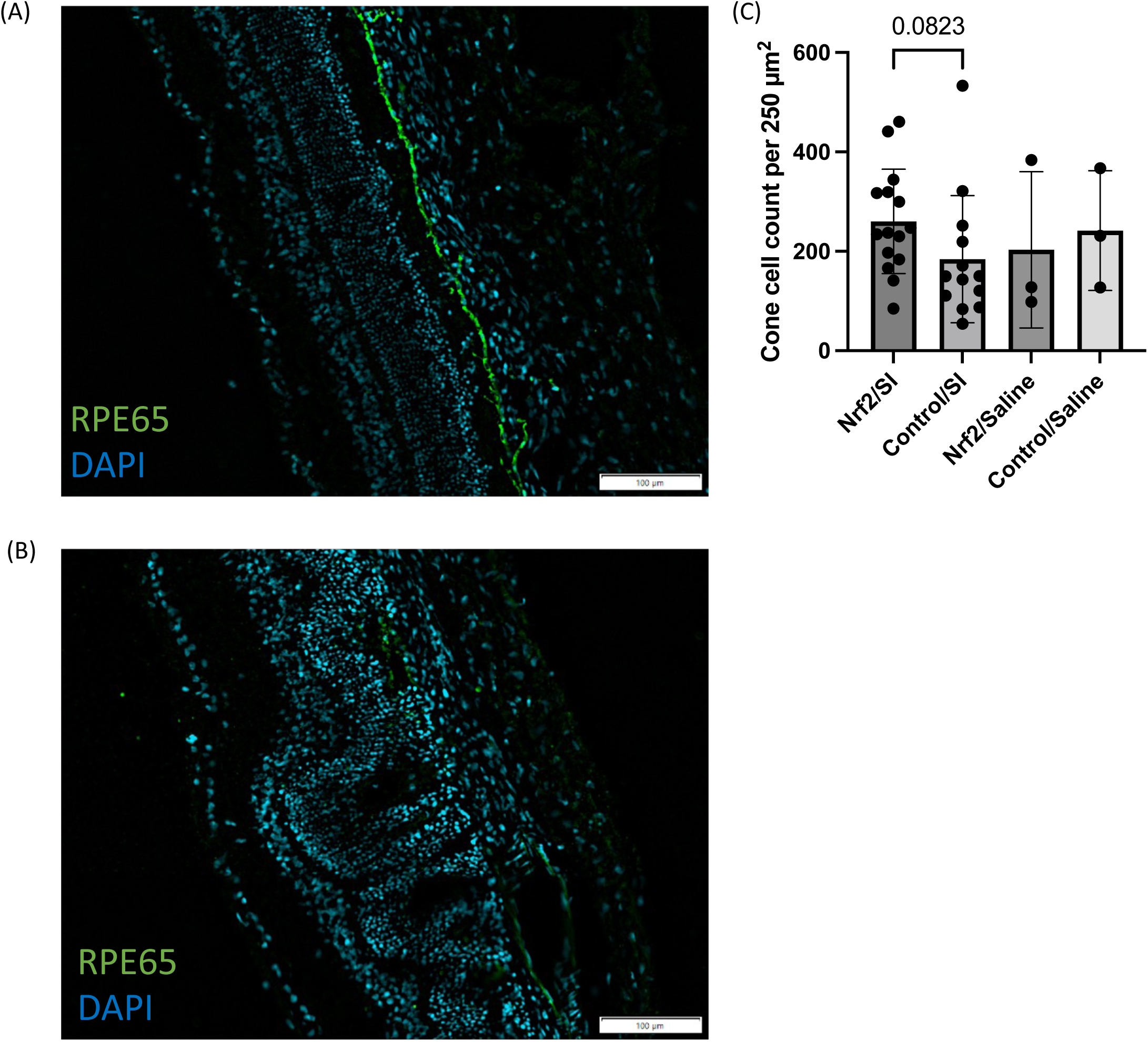
Alternative labeling/quantification strategies for assessing RPE and cone toxicity. (A) Representative eyecup sections of rat retinas (same as those in Figure 1F) 4-5 weeks following IP injection of saline. RPE65 staining is shown in green, and DAPI staining is shown in blue. Scale bar is 100 microns. (B) Same as panel (A), except an IP injection of S.I. was performed. Scale bar is 100 microns. (C) Quantification of the number of cones from rat retinal flatmounts (same as those in Figure 2C), using RedO-H2B-GFP labeling of cones as the basis for automated cone quantification in Fiji (n=15 Nrf2/S.I. flatmounts, n=13 Control/S.I. flatmounts, n=3 Nrf2/Saline flatmounts, n=3 Control/Saline eyes, mean ± SD, p=0.0823; paired T test). For this analysis, samples with median cone counts at or below 25 were excluded (indicative of partial/no transduction by AAV8/RedO-H2B-GFP following subretinal injection).

**Figure S3:**
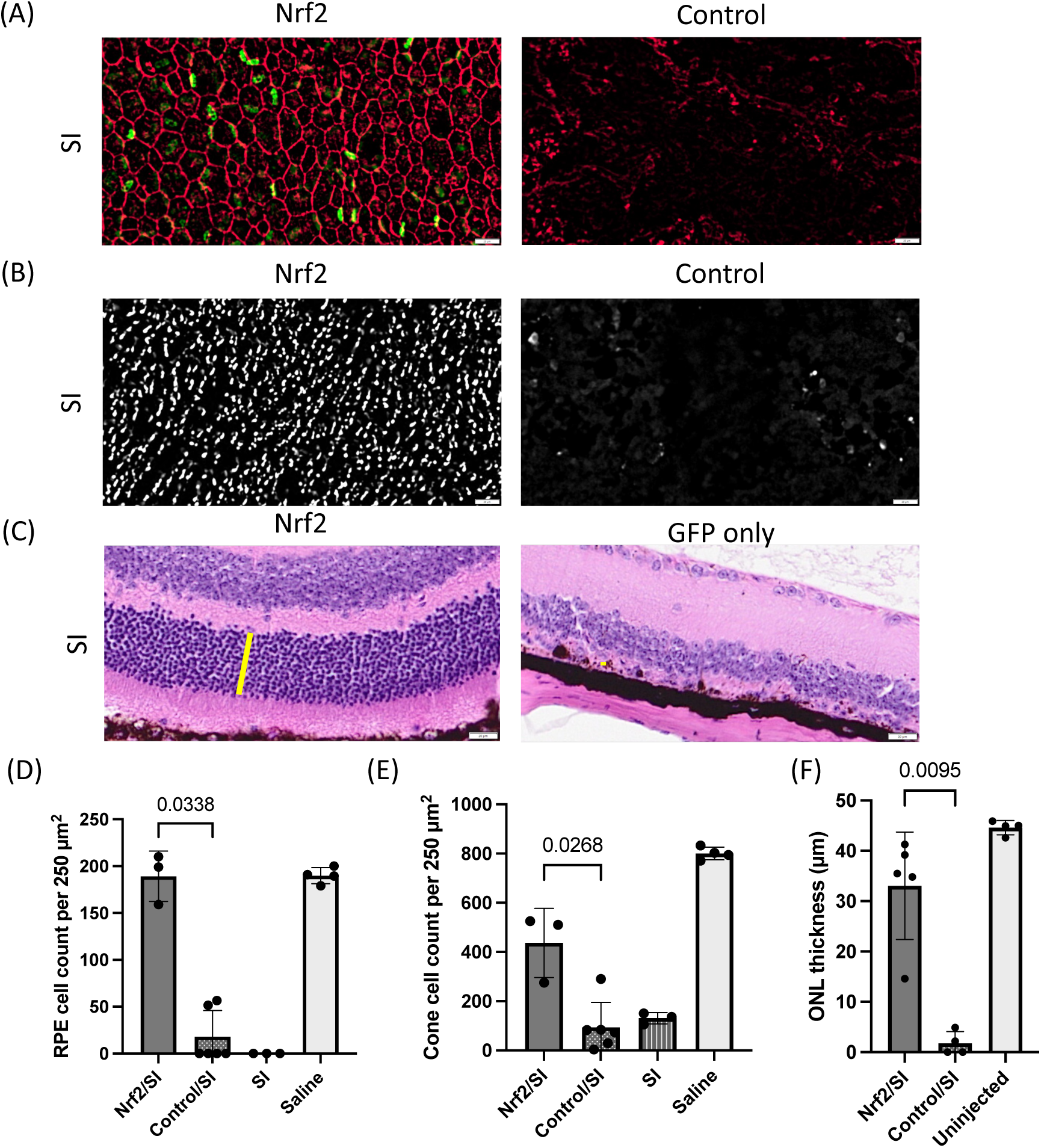
Assessment of AAV8/Best1-Nrf2 on RPE and retinal histology in mice. For panels A/B/D/E, mice were injected at birth with 4e8 vg AAV8/Best1-Nrf2 + 2e8 vg AAV8/RedO-H2B-GFP in one eye or 4e8 vg AAV8/Best1-6xSTOP-mutGFP control vector + 2e8 vg AAV8/RedO-H2B-GFP in the contralateral eye and IP injected with S.I. at 8-10 weeks of age. Tissues were harvested 12-13 weeks post IP injection. For panels C/F, mice were injected at birth with 4e8 vg AAV8/Best1-Nrf2 + 2e8 vg AAV8/RedO-H2B-GFP in one eye or 2e8 vg AAV8/RedO-H2B-GFP in the contralateral eye and IP injected with S.I. at 6-6.5 weeks of age. Eyes were harvested 12-13 weeks post IP injection and processed for hematoxylin & eosin staining. (A) Representative RPE flatmounts (central region) stained with phalloidin (red). Scale bar is 20 microns. (B) Representative retinal flatmounts (central region) stained with an antibody to CAR (white). Scale bar is 20 microns. (C) Representative eyecup sections (mid-peripheral region) stained with hematoxylin & eosin. Yellow bar indicates the ONL. Scale bar is 20 microns. (D) Quantification of the number of RPE cells (n=3 Nrf2/S.I. flatmounts, n=6 Control/S.I. flatmounts, mean ± SD, p=0.0338; paired T test). Data from n=3-4 saline and S.I. samples are replotted here from Figure S1 for reference (max age of Fig S1 references: ∼15 weeks; max age of Fig S4 samples: ∼23 weeks). (E) Quantification of the number of cones (n=3 for Nrf2/S.I. flatmounts, n=6 Control/S.I. flatmounts, mean ± SD, p=0.0268; paired T test). Data from n=3-4 saline and S.I. samples are replotted here from Figure S1 for reference (max age of Fig S1 references: ∼15 weeks; max age of Fig S4 samples: ∼23 weeks). (F) Quantification of the average ONL thickness in eyecup sections described in (C). As a healthy eye control, uninjected C57BL/6J mice (∼7-9 weeks old) were harvested and processed with hematoxylin & eosin (n=5 for Nrf2/S.I., n=4 for Control/S.I., n=4 for uninjected controls, mean ± SD, p=0.0095; paired T test).

**Figure S4.**
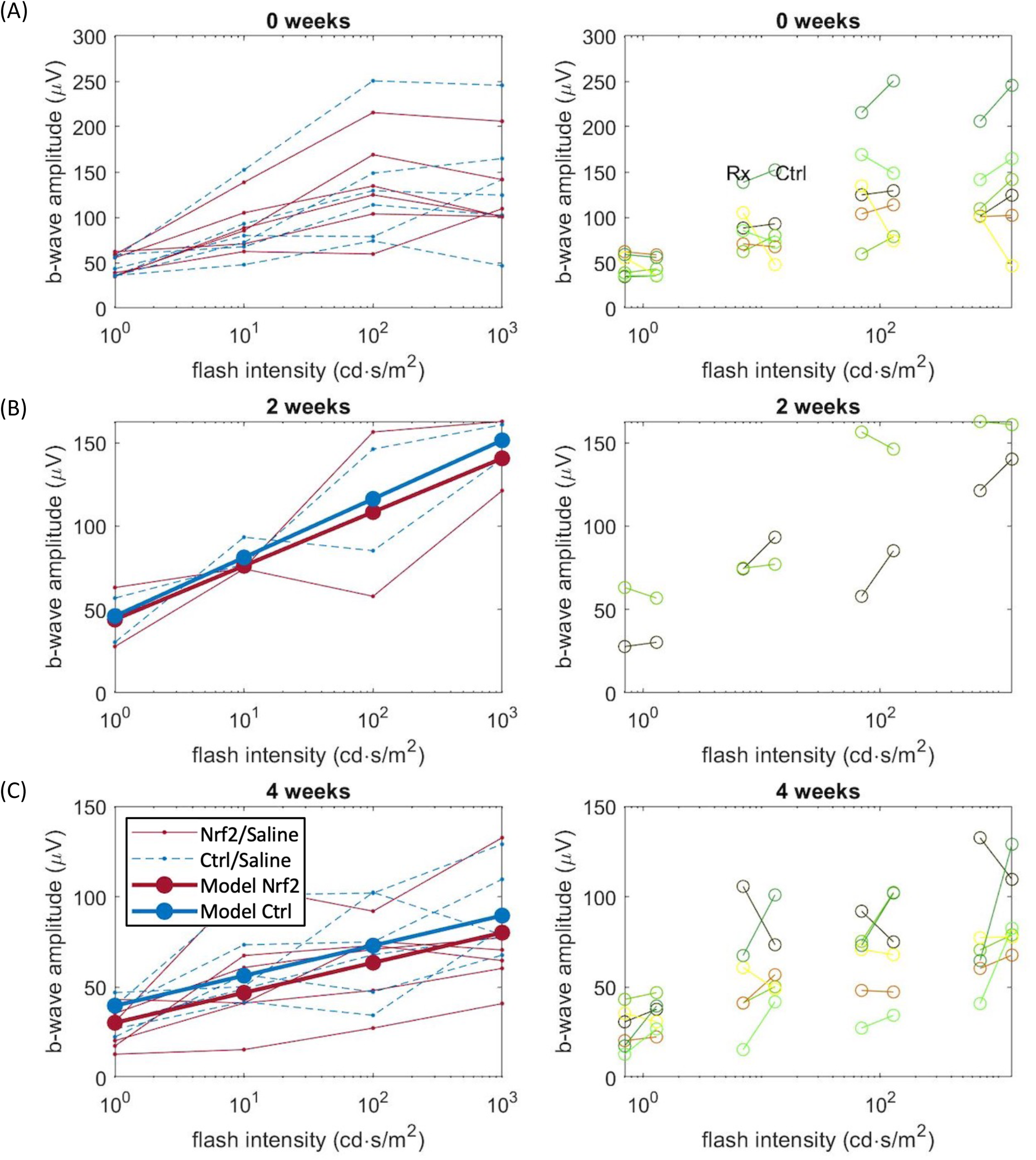
Assessment of visual function in rats injected with AAV8/Best1-Nrf2 or control AAV and IP injected with saline. Rats are from the same cohort as described in Figure 3. For all photopic ERG plots, red lines denote Nrf2/Saline treated eye data and blue lines denote control/Saline treated eye data. For the 2 week and 4 week post IP data (panels B-C), linear mixed-effects models were fit to the data using maximum likelihood with MATLAB’s ‘fitlme’ function and the model fit curves were plotted on top of the data as thicker red/blue lines (denoted Model Nrf2 and Model Ctrl in the legend). (A) Left: photopic ERG data collected before IP injection with saline (n=6 rats). Each line represents one eye of a given rat, connected across varying flash intensities. Right: the same ERG data is plotted as on the left, but lines are drawn to connect the two eyes of a given rat. (B) Same as panel A, but ERG data was collected 2 weeks post IP injection with saline (n=2 rats). (C) Same as panel A, but ERG data was collected 4 weeks post IP injection with saline (n=6 rats).

**Figure S5:**
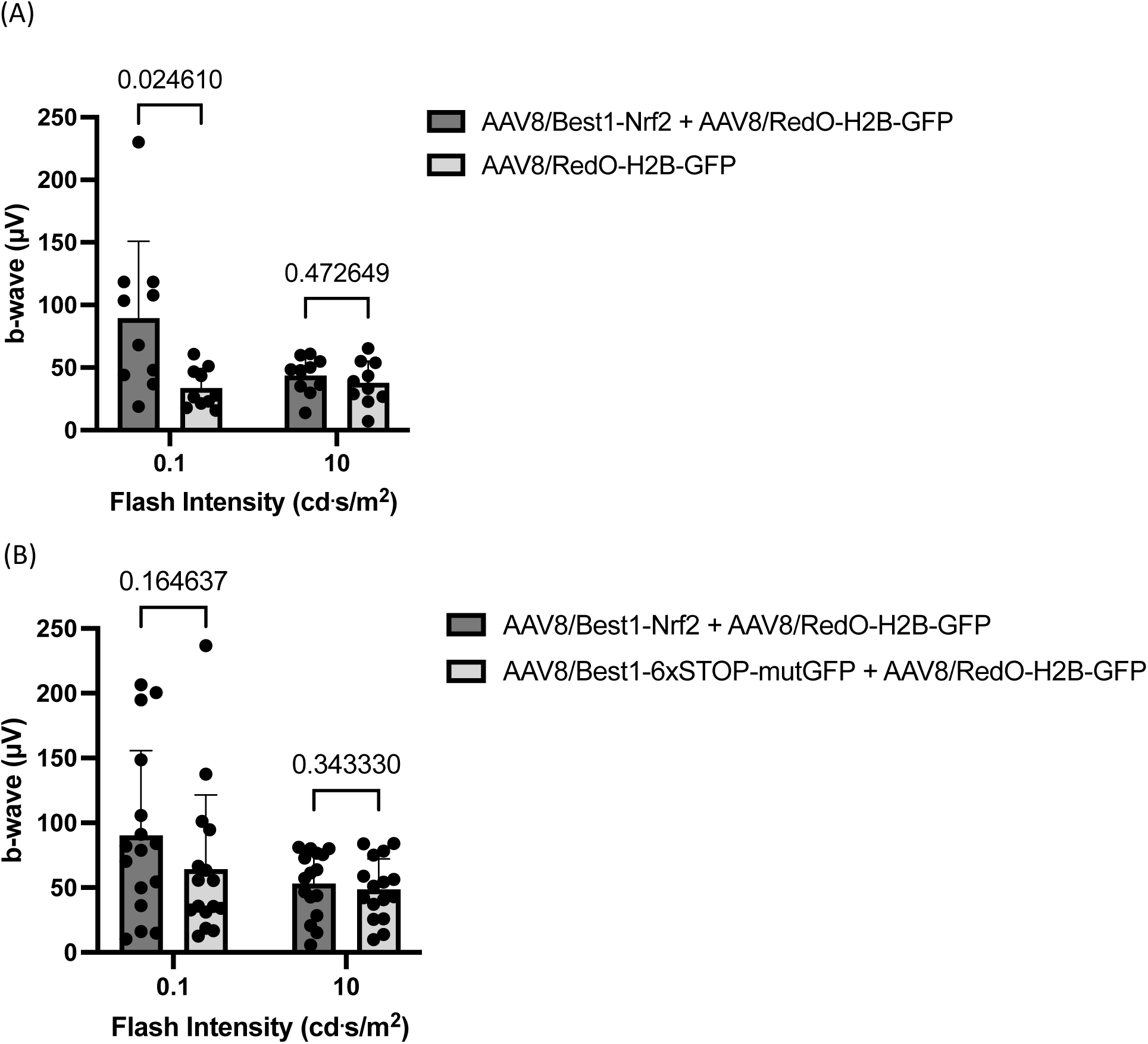
Assessment of AAV8/Best1-Nrf2 on in-life visual function in two mouse cohorts. Scotopic ERG used a flash stimulus of 0.1 cd.s/m^2^, and photopic ERG used a flash stimulus of 10 cd.s/m^2^. ERG b-wave amplitudes are plotted. All mice received an S.I. IP injection. (A) ERG data from mice injected at birth with AAV8/Best1-Nrf2 + AAV8/RedO-H2B-GFP in one eye or only AAV8/RedO-H2B-GFP in the contralateral eye and IP injected with S.I. at 6-7 weeks of age. ERG data was acquired ∼3-4 weeks post IP injection (n=10 mice, mean ± SD, p=0.0246 for scotopic, other p-values are n.s., multiple paired T tests with Bonferroni-Dunn multiple comparisons correction). (B) ERG data from mice injected at birth with AAV8/Best1-Nrf2 + AAV8/RedO-H2B-GFP in one eye or AAV8/Best1-6xSTOP-mutGFP control vector + AAV8/RedO-H2B-GFP in the contralateral eye and IP injected with S.I. at 8-10 weeks of age. ERG data was acquired ∼4-5 weeks post IP injection (n=16 mice, mean ± SD, all p-values are n.s., multiple paired T tests with Bonferroni-Dunn multiple comparisons correction).

**Figure S6:**
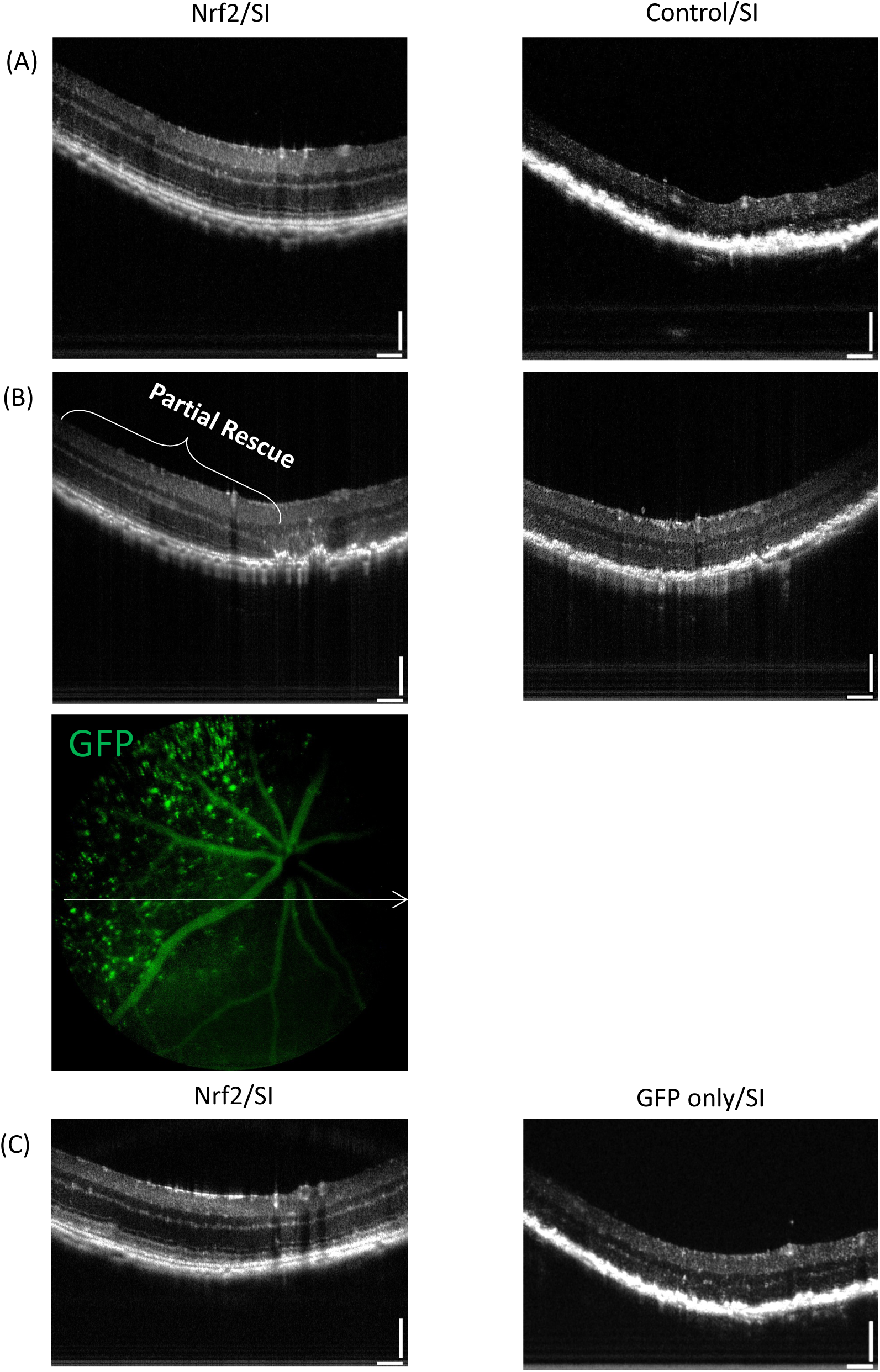
Assessment of AAV8/Best1-Nrf2 on in-life retinal structure in two mouse cohorts. (A) Representative OCT imaging in mice injected at birth with AAV8/Best1-Nrf2 + AAV8/RedO-H2B-GFP in one eye or AAV8/Best1-6xSTOP-mutGFP control vector + AAV8/RedO-H2B-GFP in the contralateral eye and IP injected with S.I. at ∼8-10 weeks of age. Images were acquired ∼4 weeks post IP injection. (B) Same as (A), except an example of partial rescue detectable by OCT and GFP (in fluorescent fundus image) is shown. (C) Same as (A), except the mice were injected at birth with AAV8/Best1-Nrf2 + AAV8/RedO-H2B-GFP in one eye or only AAV8/RedO-H2B-GFP in the contralateral eye and IP injected with S.I. at ∼6-6.5 weeks of age. Images were acquired ∼4 weeks post IP injection.

**Figure S7:**
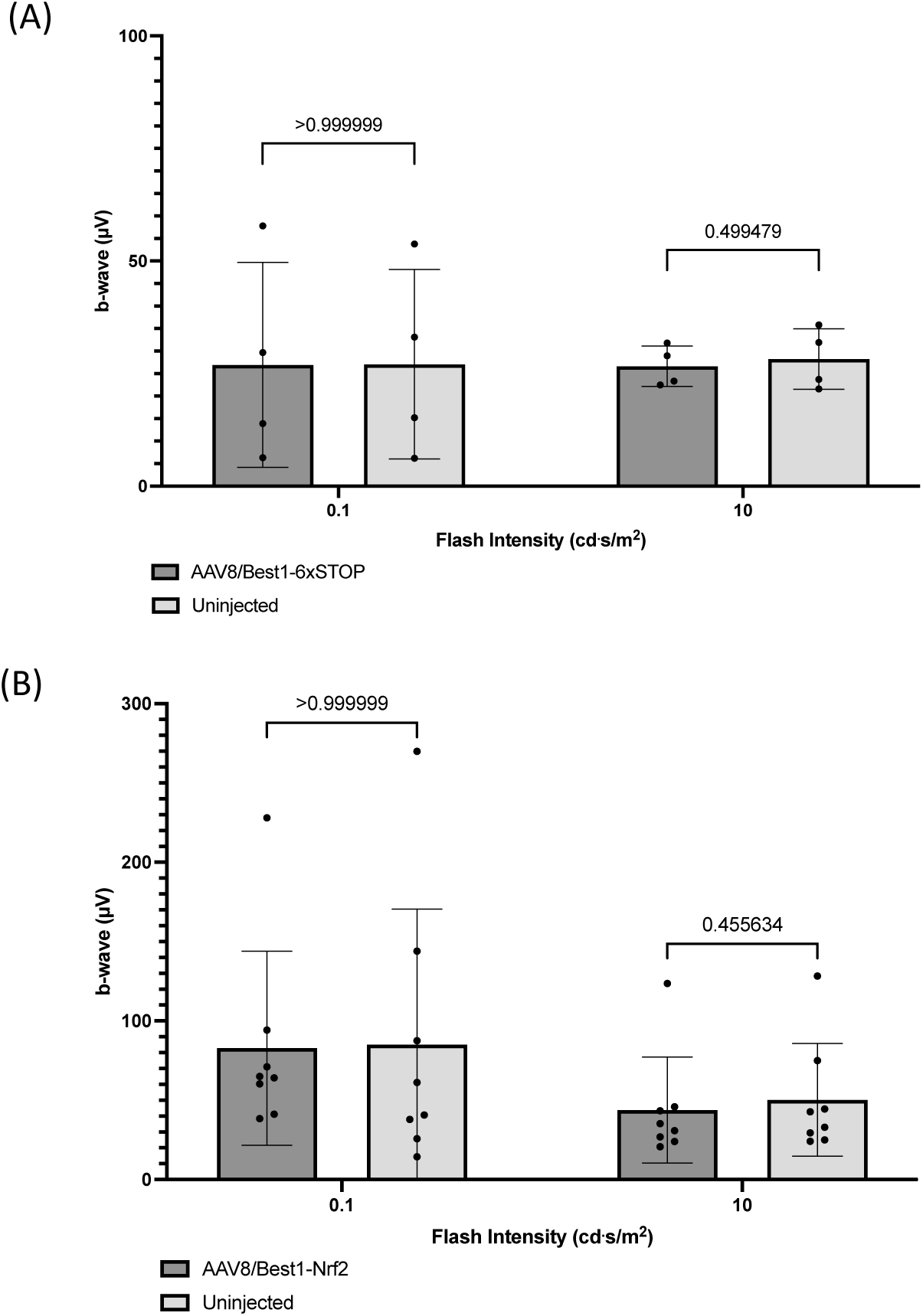
Assessment of AAV8/Best1-Nrf2 on visual function in mice subretinally injected as adults. ERG b-wave amplitudes were collected at 5 different light intensities and plotted. ERG data were collected 6-7 weeks post IP injection. All animals were IP injected with S.I. (A) ERG data from mice subretinally injected at 6-13 weeks of age with AAV8/Best1-6xSTOP control vector + AAV8/RedO-H2B-GFP in the right eye, with the left eye uninjected (n=4 mice, mean ± SD, all p-values are n.s., multiple paired T tests with Bonferroni-Dunn multiple comparisons correction). (B) ERG data from mice subretinally injected at 6-13 weeks of age with AAV8/Best1-Nrf2 + AAV8/RedO-H2B-GFP in the right eye, the left eye uninjected (n=8 mice, mean ± SD, all p-values are n.s., multiple paired T tests with Bonferroni-(A)

**Figure S8:**
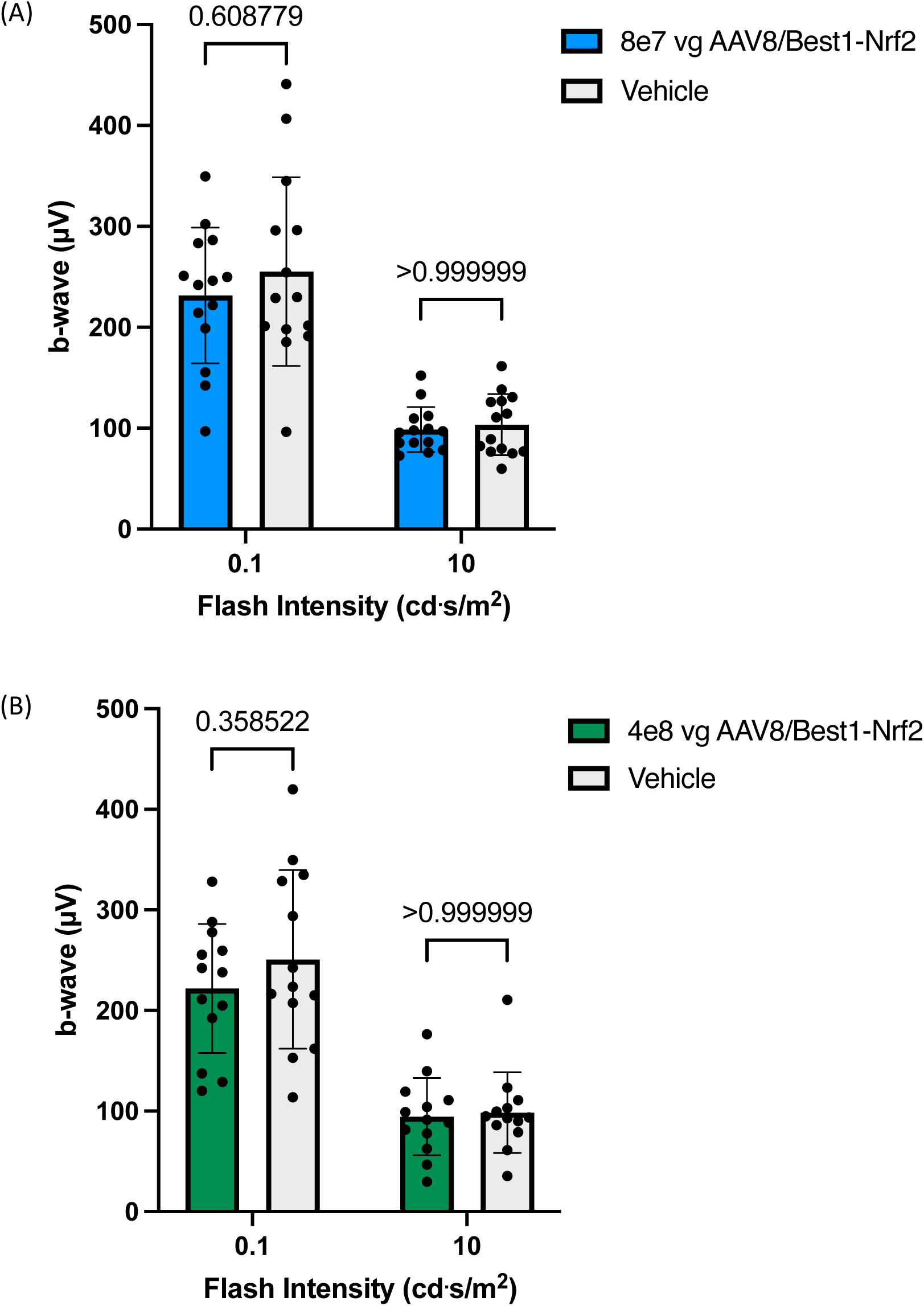
Tolerability assessments for AAV8/Best1-Nrf2 in mice. Scotopic ERG used a flash stimulus of 0.1 cd.s/m^2^, and photopic ERG used a flash stimulus of 10 cd.s/m^2^. B-wave amplitudes are plotted. ERG data was collected from mice at 1-2 months of age. (A) ERG data from mice injected with a dose of AAV8/Best1-Nrf2 (8e7 vg) in one eye and a vehicle control (see Methods) in the contralateral eye (n=14 mice, mean ± SD, all p-values are n.s., multiple paired T tests with Bonferroni-Dunn multiple comparisons correction). (B) ERG data from mice injected with a dose of AAV8/Best1-Nrf2 (4e8 vg) in one eye and a vehicle control (see Methods) in the contralateral eye (n=13 mice, mean ± SD, all p-values are n.s., multiple paired T tests with Bonferroni-Dunn multiple comparisons correction).

**Figure S9:**
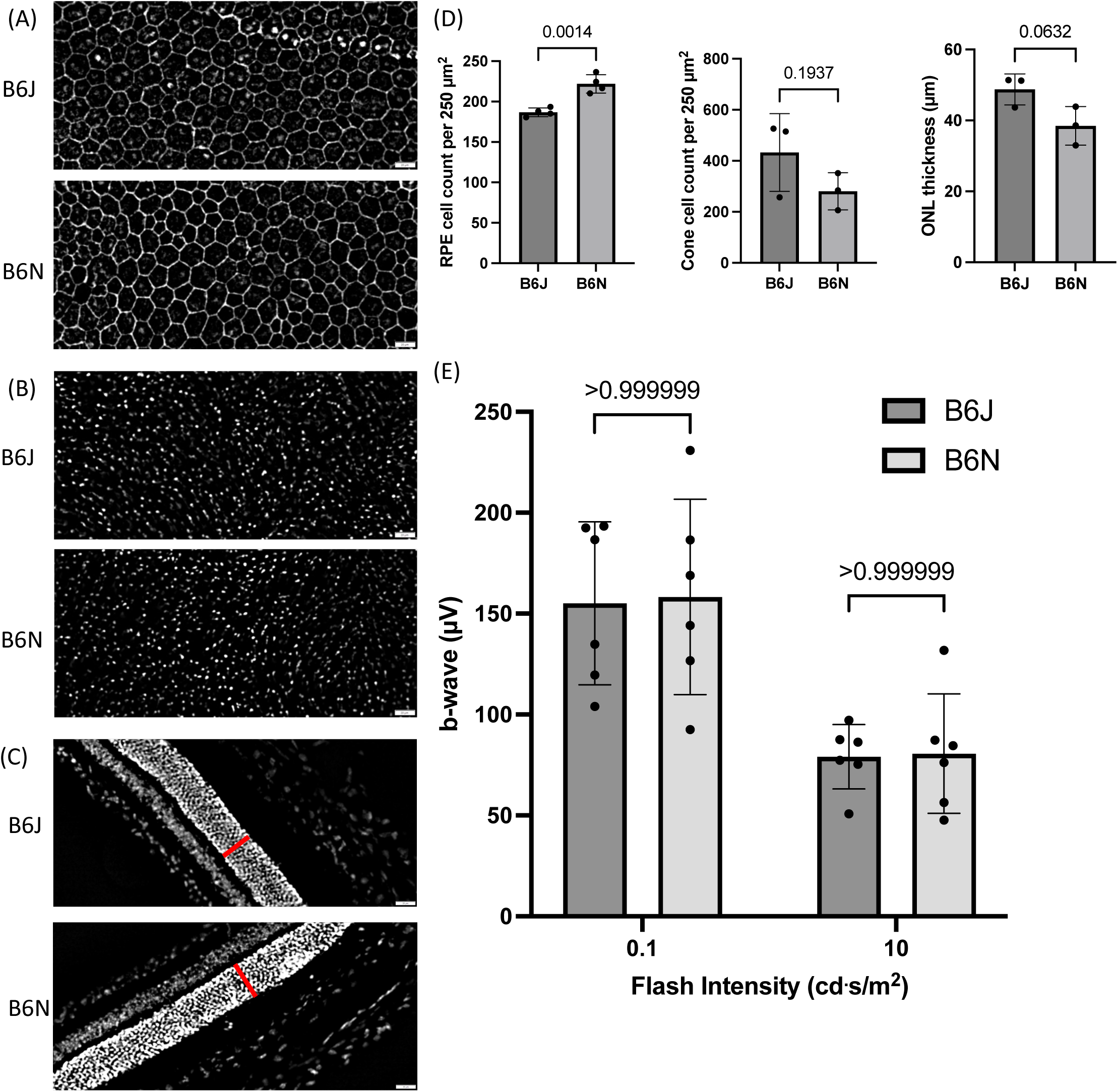
Comparison of B6J and B6N mice. C57BL/6J mice (from Jackson labs) are abbreviated B6J, and C57BL/6NCrl mice (from Charles River labs) are abbreviated B6N. B6N mice include the retinal degeneration gene, rd8. All assays were performed at 16-20 weeks of age. (A) Representative RPE flatmounts (central region). Flatmounts stained with phalloidin (white). Scale bar is 20 microns. (B) Representative retinal flatmounts (central region). Flatmounts stained with an antibody to CAR (white). Scale bar is 20 microns. (C) Representative eyecup cryosections (mid-peripheral region). Sections stained with DAPI. Scale bar is 20 microns. Red bar indicates the ONL. (D) Quantification of RPE flatmounts (n=4 mice per strain, mean ± SD, p=0.0014, unpaired T test), retinal flatmounts (n=3 mice per strain, mean ± SD, p-value is n.s., unpaired T test), or eyecup cryosections (n=3 mice per strain, mean ± SD, p-value is n.s., unpaired T test). (E) ERG b-wave amplitudes at 5 different light intensities (n=6 mice per strain, mean ± SD, all p-values are n.s., multiple unpaired T tests with Bonferroni-Dunn multiple comparisons correction).

